# Proteasome-dependent Orc6 removal from chromatin upon S-phase entry safeguards against MCM reloading and tetraploidy

**DOI:** 10.1101/2024.01.30.577900

**Authors:** Yoko Hayashi-Takanaka, Ichiro Hiratani, Tokuko Haraguchi, Yasushi Hiraoka

## Abstract

DNA replication is tightly regulated to occur only once per cell cycle, as untimely re-initiation can lead to aneuploidy, which is associated with early senescence and cancer. The pre-replication complex (comprising Orc1-6, Cdc6, Cdt1, and MCM) is essential for the initiation of DNA replication, but the dynamics and function of Orc6 during the cell cycle remain elusive. Here, we demonstrate that Orc6 associates with chromatin during G1-phase and dissociates upon S-phase entry. The dissociation of Orc6 from chromatin is dependent on proteasome activity, and inhibition of the proteasome leads to the accumulation of chromatin-bound Orc6, which promotes abnormal MCM loading after S-phase entry without undergoing mitosis in human immortalized hTERT-RPE1 cells. Following release from proteasome inhibition, cells with elevated levels of chromatin-bound Orc6 and MCM proceed to the next replication phase as tetraploid cells. Our findings suggest that the proteasome-dependent dissociation of Orc6 after DNA replication is critical for preventing inappropriate MCM reloading and tetraploid formation.

## Introduction

Genomic DNA replication is fundamental to the proliferation of living organisms. In eukaryotes, this process is tightly regulated to ensure that all chromosomes replicate only once per cell cycle. Disruptions in this regulation can result in abnormal DNA replication, leading to genomic instability, which may promote senescence or oncogenesis (Thakur et al., 2022).

DNA replication begins with the assembly of pre-replication complexes (pre-RCs) at replication origins during G1 phase (Hu and Stillman, 2023). The pre-RC consists of four key components: the origin recognition complex (ORC), minichromosome maintenance (MCM) complex, Cdc6, and Cdt1. MCM, a replicative helicase, is composed of Mcm2–7 subunits (Bell and Botchan, 2013; Deegan and Diffley, 2016; Ishimi, 2018), while Cdc6 and Cdt1 facilitate the recruitment of MCM to chromatin. Upon entry into S-phase, the pre-RC disassembles. Aberrant loading of pre-RCs onto chromatin after the S-phase initiation can trigger DNA re-replication without mitosis (Vassilev and DePamphilis, 2017; Limas and Cook, 2019), underscoring the need for tight regulation of Cdt1 and Cdc6 levels by proteasome degradation through E3 ubiquitin ligases to prevent pre-RC formation beyond G1 phase (Nishitani et al., 2006; Walter et al., 2016).

The ORC is a six-subunit complex (Orc1–6), which binds chromatin to mark replication origins for pre-RC assembly during G1 phase (Bell and Labib, 2016; Ganier et al., 2019; Hu and Stillman, 2023). While Orc2–5 remain chromatin-bound throughout the cell cycle, Orc1 dissociates from chromatin after S-phase entry via ubiquitin-mediated proteolysis (Li and DePamphilis, 2002; Méndez et al., 2002; Tatsumi et al., 2003; Coulombe et al., 2019). Orc1–5 share structural similarities, whereas Orc6 is more similar to TFIIB (Duncker et al., 2009; Liu et al., 2011; Feng et al., 2021), distinguishing it functionally from other ORCs. Orc6 can bind DNA independently of Orc1-5 (Xu et al., 2020) and has roles in chromosome segregation and cytokinesis (Prasanth et al., 2002; Chesnokov et al., 2003; Semple et al., 2006; Balasov et al., 2009; Bernal and Venkitaraman, 2011). Recent studies suggest additional functions for Orc6, including its involvement in replication fork progression, ATR activation, and mismatch repair (Lin et al., 2022). However, the specific role of human Orc6 in the cell cycle remains largely unexplored.

Proteasomes are the primary machinery for protein degradation in cells. Reduced proteasome activity is observed in replicative senescent cells (Sabath et al., 2020) and inhibiting proteasomes induces cellular senescence in normal human fibroblasts (Chondrogianni & Gonos, 2004; Chondrogianni et al., 2003; Torres et al., 2006; Ukekawa et al., 2004). Senescent cells, which bypass mitosis, exhibit chromatin-bound pre-RCs, suggesting a predisposition to enter S-phase as tetraploid cells (Johmura et al., 2014; Panopoulos et al., 2014; Zeng et al., 2023). However, it remains unclear whether reduced proteasome activity in cellular senescence is involved in pre-RC formation.

In this study, we explored the dynamics of Orc6 during the cell cycle in human cells using multicolor immunofluorescence-based single-cell plot analysis (hereafter referred to as “single-cell plot analysis”). This method allows for the quantitative measurements of Orc6 levels in individual cells from fluorescence microscopy images, eliminating the need for cell synchronization (Hayashi-Takanaka et al., 2020; Hayashi-Takanaka et al., 2021). Our findings suggest that the regulation of Orc6 localization throughout the cell cycle is crucial for proper pre-RC formation, and its dysregulation can lead to replication as tetraploid cells.

## Results

### Dissociation of Orc6 from chromatin during S-phase

To quantify chromatin-bound Orc6 in individual human cells, we performed single-cell plot analysis using fluorescence microscopy to measure multiple intracellular components in each cell (Hayashi-Takanaka et al., 2020; Hayashi-Takanaka et al., 2021). An asynchronous population of hTERT-RPE1 cells was fixed using two different methods: “direct fixation” and “pre-extraction” (Fig. 1A). The direct fixation method detects whole-cell proteins, while the pre-extraction method identifies only the chromatin-bound fraction (Hayashi-Takanaka et al., 2021). The fixed cells were co-stained with Hoechst 33342 (a DNA indicator, referred to as “Hoechst”), EdU (an S-phase marker), and an anti-Orc6 antibody (Fig. 1A). The same cells were also stained with an anti-Orc2 antibody to detect the behavior of Orc2–5 subunits on chromatin throughout the cell cycle. The specificity of these antibodies was validated by immunostaining as shown in Fig. S1.

**Figure 1.**
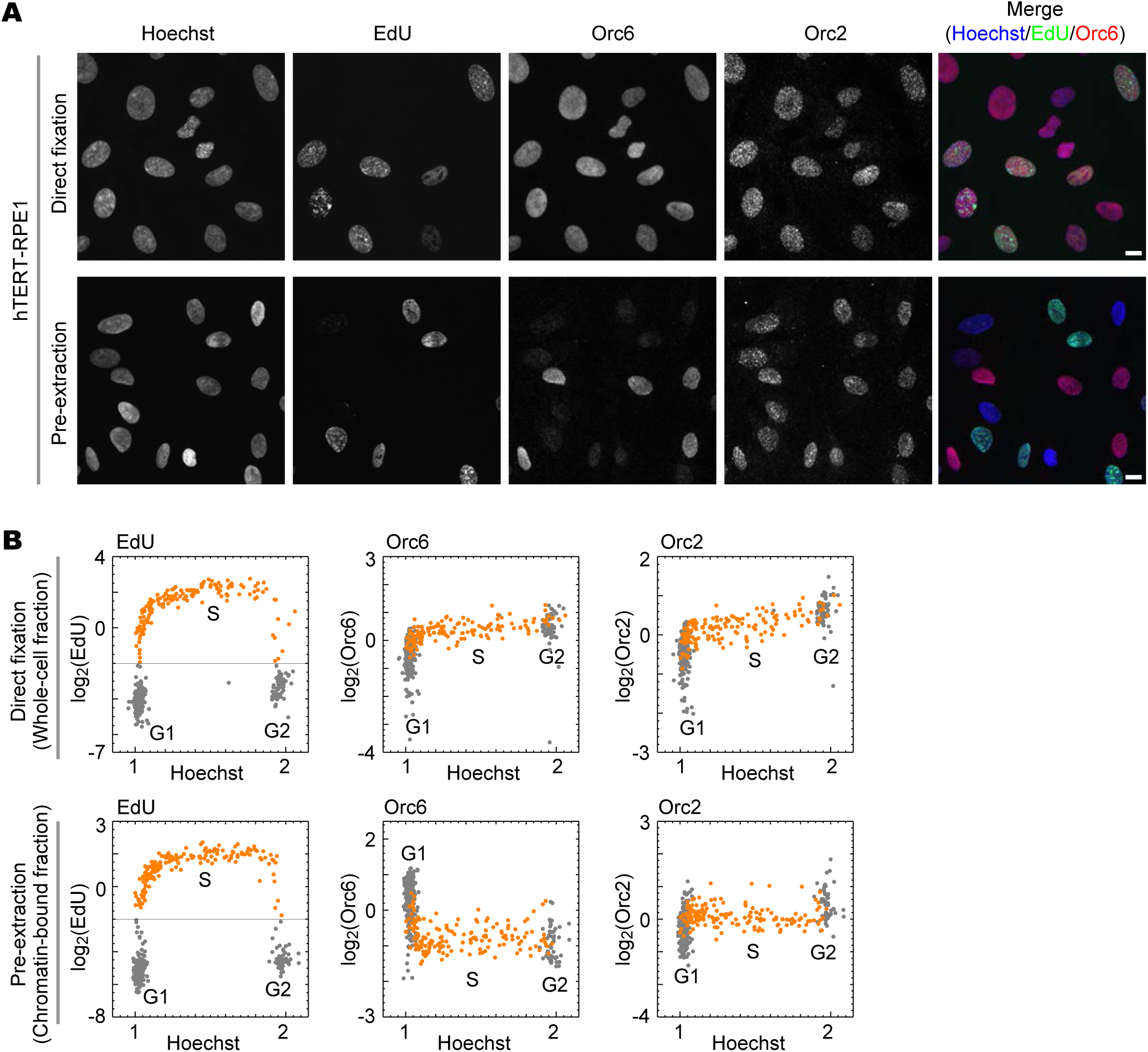
Detection of chromatin-bound Orc6 by single-cell plot analysis. (**A**) Representative fluorescence images of Hoechst, EdU, Orc6, Orc2, and merged (Hoechst/EdU/Orc6) in hTERT-RPE1 cells prepared using the direct fixation method (upper panels) and the pre-extraction method (lower panels). Scale bars, 10 µm. (**B**) Single-cell plot analysis based on the images in (**A**). Each dot represents the EdU, Orc6, and Orc2 intensities in a single individual cell against the Hoechst intensities. The orange dots represent S-phase cells based on EdU intensities (log_2_[EdU] > ­2). n = 450 (number of cells examined in each panel).

Fluorescence imaging revealed that Orc6 staining was relatively uniform in the cell population when using the direct fixation method but showed variability among the cells prepared by the pre-extraction method (compare upper and lower panels for Orc6 in Fig. 1A). In contrast, Orc2 staining appeared punctate and remained consistent across all cells in both methods (right panels, Fig. 1A). Single-cell plot analysis was performed based on these images: the DNA content was determined by Hoechst intensity normalized from 1 to 2, and the normalized intensities of EdU, Orc6, and Orc2 were plotted against the DNA content, where each dot represents a single cell (Fig. 1B). EdU-positive S-phase cells (>­2 on the log_2_ scale), marked as orange dots in the EdU panels (left panels, Fig. 1B), are also shown as orange dots in both the Orc6 and Orc2 panels (middle and right panels, Fig. 1B). In the direct fixation method, the whole-cell fraction of Orc6 levels remained relatively constant, ranging from ­1 to 1 on the log_2_ scale throughout the cell cycle (upper middle panel, Fig. 1B). However, using the pre-extraction method, chromatin-bound Orc6 levels were high during G1-phase, dropped immediately after S-phase entry, and stayed low through S- and G2-phases (lower middle panel, Fig. 1B). Orc2 levels remained relatively constant during the G1/S transition and showed minimal changes during the S-phase in both fixation methods (right panels, Fig. 1B). These results indicate that Orc6, but not Orc2, dissociates from chromatin during S-phase in hTERT-RPE1 cells.

To determine if Orc6 dissociation occurs in other cell types, we analyzed changes in chromatin-bound Orc6 levels during the cell cycle in cancer cell lines (HeLa and U2OS cells) and normal fibroblast cells (IMR90 cells) using the pre-extraction method (Fig. S2A). The reduction in Orc6 levels during the G1- to S-phase transition was less pronounced in cancer cell lines (HeLa and U2OS cells, Fig. S2B) compared to normal cell lines (IMR90 and hTERT-RPE1 cells, Fig. S2B). This suggests that the regulation of chromatin-bound Orc6 levels during the cell cycle may differ between normal and cancer cells, or among different cell types.

### Involvement of the proteasome in Orc6 dissociation from chromatin

Since the levels of Cdt1, a component of the pre-RC, are regulated by proteasome activity (Nishitani et al., 2006; Walter et al., 2016), we examined whether Orc6 is also a target for proteasome regulation using the proteasome inhibitor MG132. hTERT-RPE1 cells were treated with 20 µM MG132 for varying durations (1, 2, and 4 h), fixed using two different fixation methods, and stained with Hoechst and an anti-Orc6 antibody. The same cells were co-stained with anti-Cdt1 and anti-Orc2 antibodies as controls for proteasome degradation (Fig. 2A). Single-cell plot analysis revealed that Orc6 levels in the whole-cell fraction (obtained by the direct-fixation method) remained relatively unchanged during MG132 treatment (blue dots, left panels in Fig. 2A). However, Orc6 levels in the chromatin-bound fraction (obtained by the pre-extraction method) increased during MG132 treatment (blue dots, right panels in Fig. 2A). Without MG132 treatment, Orc6 was predominantly bound to chromatin during G1-phase and dissociated after entering S-phase (0 hr, Fig. 2A). Upon the treatment with MG132 for 1 h, the decrease in Orc6 levels at the G1- to S-phase transition was less pronounced compared to 0 hr. After 2 h of MG132 treatment, Orc6 remained bound to chromatin after the S-phase, similar to its levels in G1-phase. As a control, Cdt1 levels increased in cells progressing through S and G2 phases after 1 h of treatment and continued to increase with extended treatment in both whole-cell fraction and chromatin-bound fraction (orange dots, Fig. 2A), indicating that proteasome degradation was functioning normally under these experimental conditions. Orc2 levels remained relatively unchanged for up to 4 h in the whole-cell fraction (green dots, left panels in Fig. 2A), and slightly increased after 4 h in the chromatin-bound fraction (green dots, right panels in Fig. 2A). These results suggest that Orc6, and to a lesser extent Orc2, dissociates from chromatin via proteasome activity during the S and G2 phases, but is not degraded by the proteasome.

**Figure 2.**
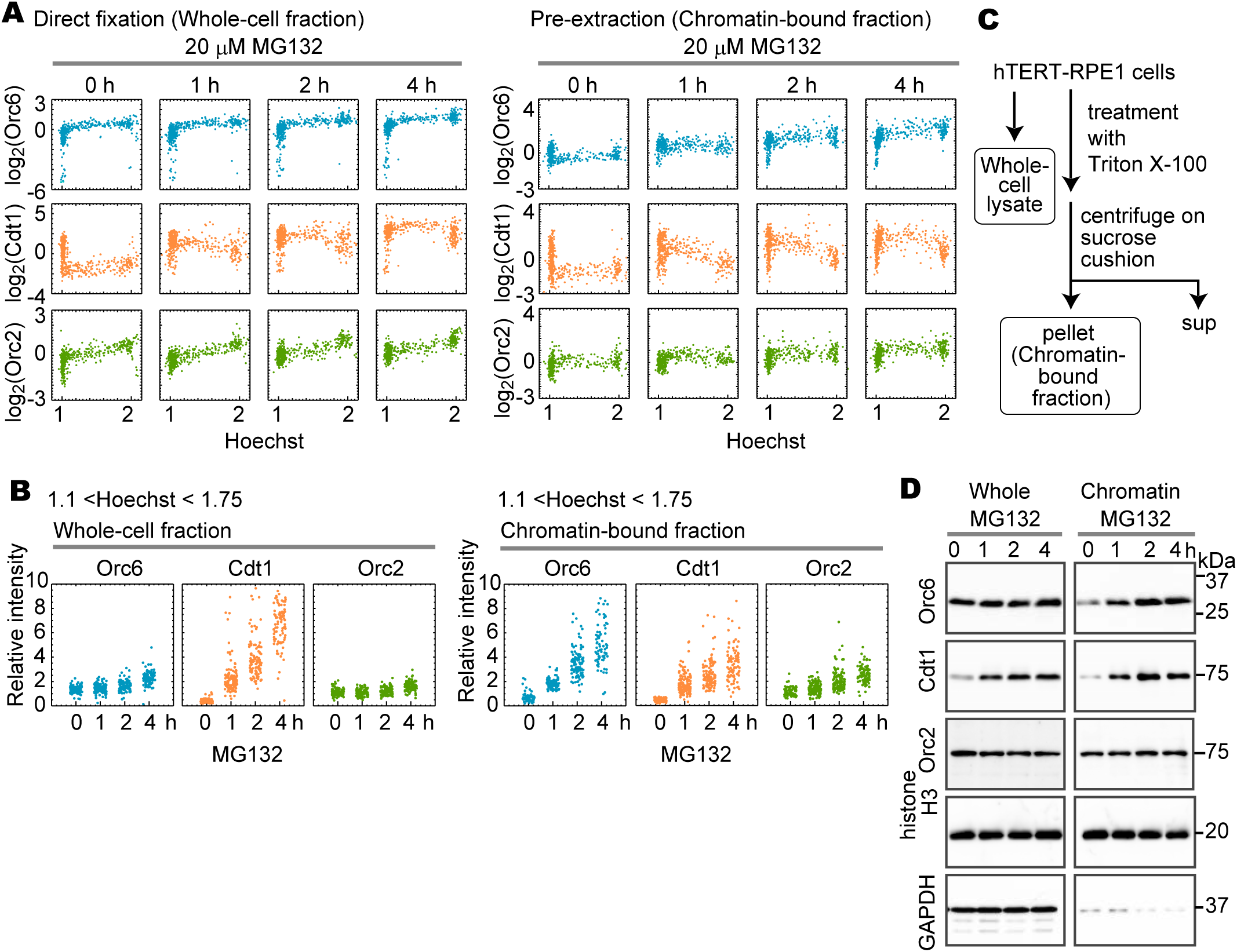
Effects of proteasome inhibitors on the dynamics of Orc6 during the cell cycle. Single-cell plot analysis of Orc6 (top), Cdt1 (middle), and Orc2 (bottom) in hTERT-RPE1 cells. Cells were treated with 20 µM MG132 for 0, 1, 2, or 4 h (timescale on the top), fixed using the direct fixation method (left panels) and the pre-extraction method (right panels). Each dot represents the intensity of Orc6, Cdt1, and Orc2 in a single individual cell plotted against the Hoechst intensity. n = 450 (number of cells examined in each panel). (**B**) Orc6, Cdt1, and Orc2 levels in the S-phase cells. Cells corresponding to the S-phase (1.1 < Hoechst < 1.75) were selected from (**A**), and their fluorescence intensities were plotted. The average intensity in MG132-untreated cells (0 h) was defined as 1. G1- and G2-phase cells are shown in Fig. S3A, B. (**C**) Schematic diagram of the biochemical fractionation method for whole-cell lysates and chromatin-bound fractions. (**D**) Western blot analysis using antibodies against Orc6, Cdt1, Orc2, histone H3, and GAPDH. hTERT-RPE1 cells were treated with 20 μM MG132 for the indicated periods (h). Whole-cell lysates (left) and chromatin-bound fractions (right) were prepared and analyzed using western blotting. Histone H3 and GAPDH are loading controls. Molecular weight markers (kDa) are indicated on the right.

To compare the increase in Orc6 levels during the S-phase due to MG132 treatment with that of Cdt1 and Orc2, S-phase cells were selected based on Hoechst intensity (1.1< Hoechst <1.75) from the single-cell plot analysis (Fig. 2A), and then the fluorescence intensity was plotted during the time course of MG132 treatment (each dot represents a single cell, Fig. 2B). The same analysis was carried out also in G1-phase cells (Hoechst <1.1; Fig. S3A) and G2-phase cells (1.75< Hoechst; Fig. S3B). The results showed little to no increase in Orc6 and Orc2 levels in the whole-cell fraction after up to 4 h of treatment (1–2-fold; whole-cell fraction, Fig. 2B), while a large increase in Cdt1 levels was observed with increasing duration of MG132 treatment (up to 8-fold; left panels, Fig. 2B and Fig. S3A, B). In the chromatin-bound fraction, Orc6 levels in S-phase cells showed a substantial increase with prolonged MG132 treatment (up to 8-fold; right panels, Fig. 2B), while those in G1- and G2-phase cells increased mildly (up to 4-fold; right panels, Fig. S3A, B). The increase in chromatin-bound Cdt1 and Orc2 levels was smaller compared to Orc6 (up to 5-fold and 3-fold, respectively; chromatin-bound fraction, Fig. 2B). The MG132-induced increase in chromatin-bound Orc6 was independent of the presence of Orc1 or Orc2 because Orc6 binding to chromatin was not affected by the siRNA depletion of either Orc1 or Orc2 as shown later in Fig. S6A, B and D.

To further confirm these results, we examined Orc6 protein levels in whole-cell lysates and chromatin-bound fractions obtained via sucrose cushion fractionation (Fig. 2C). Cells were treated with 20 µM MG132 for varying durations (1, 2, and 4 h), and Orc6 levels were analyzed in whole-cell lysates and chromatin-bound fractions by western blot analysis and quantified in three independent experiments (Fig. S3C). Orc6 levels in whole-cell lysates remained unchanged throughout MG132 treatment (Fig. 2D, Fig. S3C). In contrast, the levels of Orc6 in the chromatin-bound fractions increased approximately 2-fold (Fig. 2D, Fig. S3C). After treatment with MG132 for 2–4 hours, the chromatin-bound fraction of Orc6 was nearly equivalent to the whole-cell fraction (Fig. 2D, Fig. S3C, Supplementary information “Blot transparency”), indicating that most Orc6 binds to chromatin upon the inhibition of proteasome activity. The amount of Cdt1 was also examined as a control to confirm the effect of the proteasome inhibitor (Fig. 2D). Cdt1 levels increased over time in both whole-cell lysates (approximately 7-fold) and chromatin-bound fractions (approximately 3-fold) after MG132 treatment. Orc2 levels remained relatively unchanged, with only a slight increase (1.2–1.5-fold) (Fig. 2D). These results are consistent with the single-plot analysis (Fig. 2A, B, S3A, B), suggesting that Orc6 dissociates from chromatin during the S-phase via proteasome activity, without undergoing degradation.

### Abnormal chromatin loading of Orc6 occurs in aged cells

In our analysis of Orc6 protein levels across various cell types (Fig. S2), we observed that certain IMR90 cells (human diploid fibroblasts) exhibited abnormally high levels of chromatin-bound Orc6 within enlarged nuclei (Fig. 3A). Since such enlarged nuclei often appear in aged cells, we hypothesized that the aging of IMR90 cells accounts for the high Orc6 levels. To test this hypothesis, we examined the levels of chromatin-bound Orc6 and MCM during the cell cycle in IMR90 cells from passage 18 and passage 32. Cells were fixed using the pre-extraction method and stained with Hoechst, anti-Orc6, and anti-Mcm2 antibodies (Fig. 3A). Single-cell plot analysis showed that, in passage 18 cells, the dynamics of chromatin-bound Orc6 and Mcm2 throughout the cell cycle were similar to those observed in hTERT-RPE1 cells. Specifically, chromatin-bound Mcm2 levels increased during G1 phase, peaked at the G1/S transition, and subsequently decreased during S phase, reaching their lowest levels in G2 phase (upper right panel in Fig. 3B, Fig. S4B) (Håland et al., 2015; Hayashi-Takanaka et al., 2021; Zeng et al., 2023; Zhou et al., 2020). Chromatin-bound Orc6 levels were high in G1 phase and decreased during S/G2 phase, similar to those seen in hTERT-RPE1 cells (upper left panel in Fig. 3B, second right in Fig. S2B). However, in passage 32 cells, a subset of cells displayed abnormally high levels of Orc6 (Orc6 >0.85 on the log2 scale) after the completion of DNA replication (1.8 < Hoechst < 2.1, shown as magenta dots in the Orc6 panel, lower left panel in Fig. 3B). When these cells with high Orc6 levels (magenta dots) were plotted on the Mcm2 panel, most of them exhibited high Mcm2 levels (red-boxed region, Fig. 3B) instead of the expected low levels (blue-boxed region, Fig. 3B). This suggests that, in passage 32 IMR90 cells, pre-RC formation, which normally occurs in G1 phase, was abnormally retained in G2 phase. These cells progressed further through the cell cycle, leading to the formation of cells with large nuclei (2 < Hoechst < 4 in lower panels, Fig. 3B). Such cells were rarely observed in passage 18 (upper left panel in Fig. 3B). These cells were characterized by SA-βGal (senescence-associated β-galactosidase), a hallmark of cellular senescence, and p21, a cell cycle inhibitor. In passage 18 cells, low levels of SA-βGal and p21 protein were observed (Fig. S4A). In contrast, cells from passage 32, particularly those with large nuclei, showed high levels of SA-βGal and/or p21 protein (Fig. S4A), confirming that IMR90 cells undergo aging between passages 18 and 32. Consequently, cells with high levels of chromatin-bound Orc6 and MCM, forming pre-RC, were more frequently observed in repeatedly passaged cells than in cells at earlier passages.

**Figure 3.**
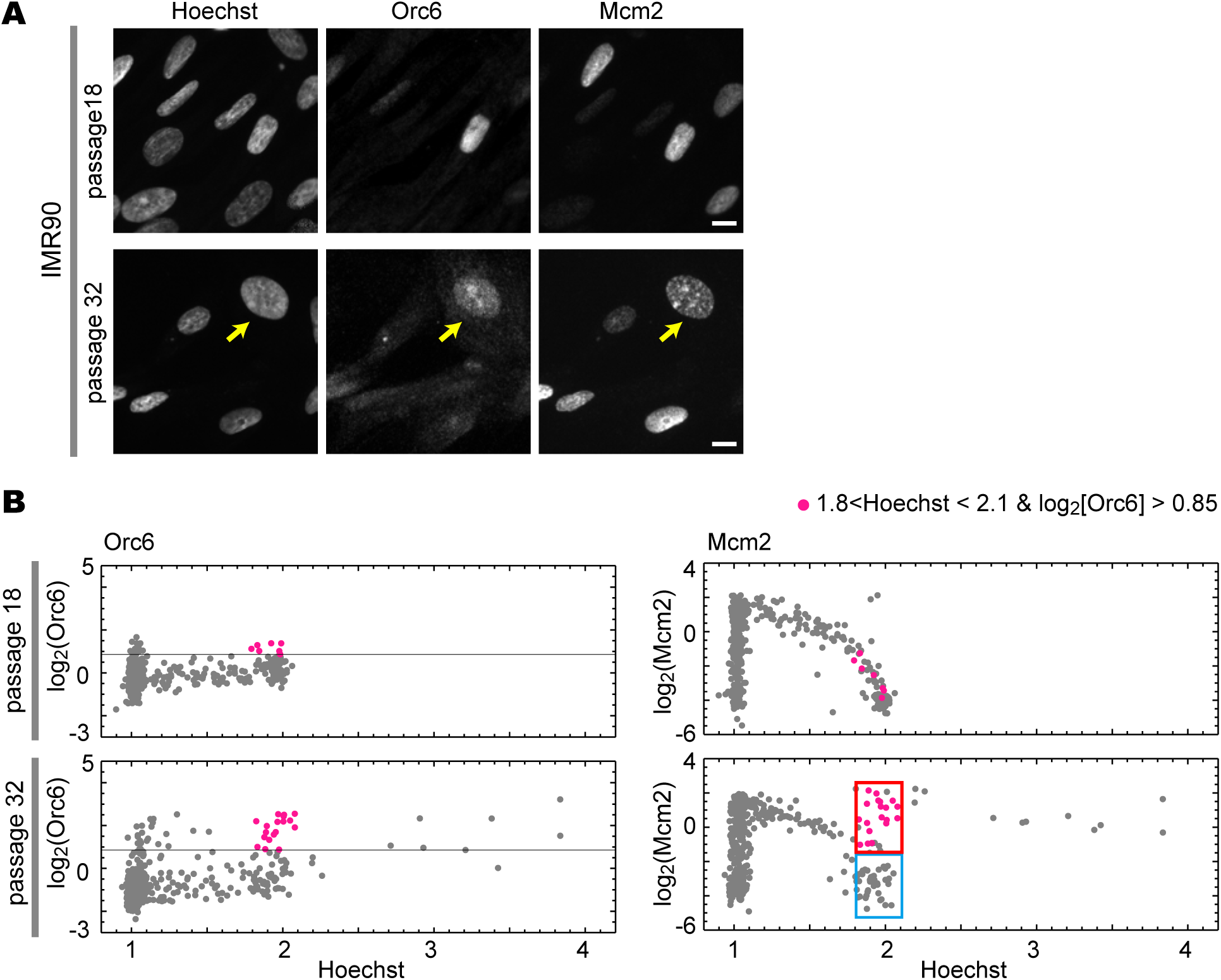
Aberrantly high levels of chromatin-bound Orc6 and Mcm2 in aged fibroblast cells. (**A**) IMR90 cells (passage numbers 18 and 32) were fixed using the pre-extraction method and stained using Hoechst, anti-Orc6, and anti-Mcm2 antibodies. Representative fluorescence images are shown. Arrows indicate relatively larger nuclei with high Orc6 signals. Scale bars, 10 µm. (**B**) Single-cell plot analysis based on the images in (**A**). Each dot represents the intensity of Orc6 and Mcm2 in an individual cell plotted against Hoechst intensity. In the range of 1.8 < Hoechst < 2.1, the magenta dots in the Orc6 panels represent cells with fluorescent signals of log_2_(Orc6) > 0.85. The same cells are also marked magenta in the Mcm2 panels. The red-boxed regions represent cells with high MCM levels (log_2_[Mcm2] > ­1.5), and the blue-boxed regions represent cells with low MCM levels (log_2_[Mcm2] < ­1.5) in the range of 1.8 < Hoechst < 2.1. n = 450 (number of cells examined in each panel).

### Abnormal chromatin loading of Orc6 occurs under low proteasome activity

As impaired proteasome function is known to occur in aged cells (Chondrogianni et al., 2003; Sabath et al., 2020), we hypothesized that the emergence of cells with abnormal loading of Orc6 and MCM in aged cells is related to reduced proteasome activity. To test this hypothesis, we examined whether reduced proteasome activity in immortalized cells (hTERT-RPE1 cells) leads to the emergence of cells with high levels of Orc6 and MCM loading on chromatin after replication, similar to the results observed in passage 32 IMR90 cells. To determine the concentrations of MG132 that reproduce cellular senescence, hTERT-RPE1 cells were treated with various concentrations of MG132 for longer periods (18 hours). Whereas most cells died at high concentrations used in Fig. 2 (> 625 nM), cell death was less frequent at a lower concentration of 312 nM MG132. Thus, we used MG132 at 312 nM or lower concentrations in the following experiments.

As a marker of cellular senescence, we examined the levels of p21 that increases upon MG132 treatment with p53 stabilization (Kwon et al., 2002; Zhu et al., 2007). Western blot analysis and immunostaining revealed elevated p21 protein levels in MG132-treated cells compared to the DMSO control (Fig. 4A, Fig. S4C), suggesting that proteasome inhibition by MG132 induces cell cycle arrest. Similarly, cells positive for SA-βGal were not detected on day 2 of treatment but appeared on day 6 (Fig. S4D). Conversely, Ki67, a protein expressed in all phases of the cell cycle except G0 phase (Scholzen and Gerdes, 2000), was widespread in MG132-treated cells on day 1 but had disappeared in some cells on day 4 (Fig. S4E). Therefore, treatment with low concentrations of MG132 for 18-24 h primarily led to cell cycle arrest, while continuous treatment over several days induced cellular senescence.

**Figure 4.**
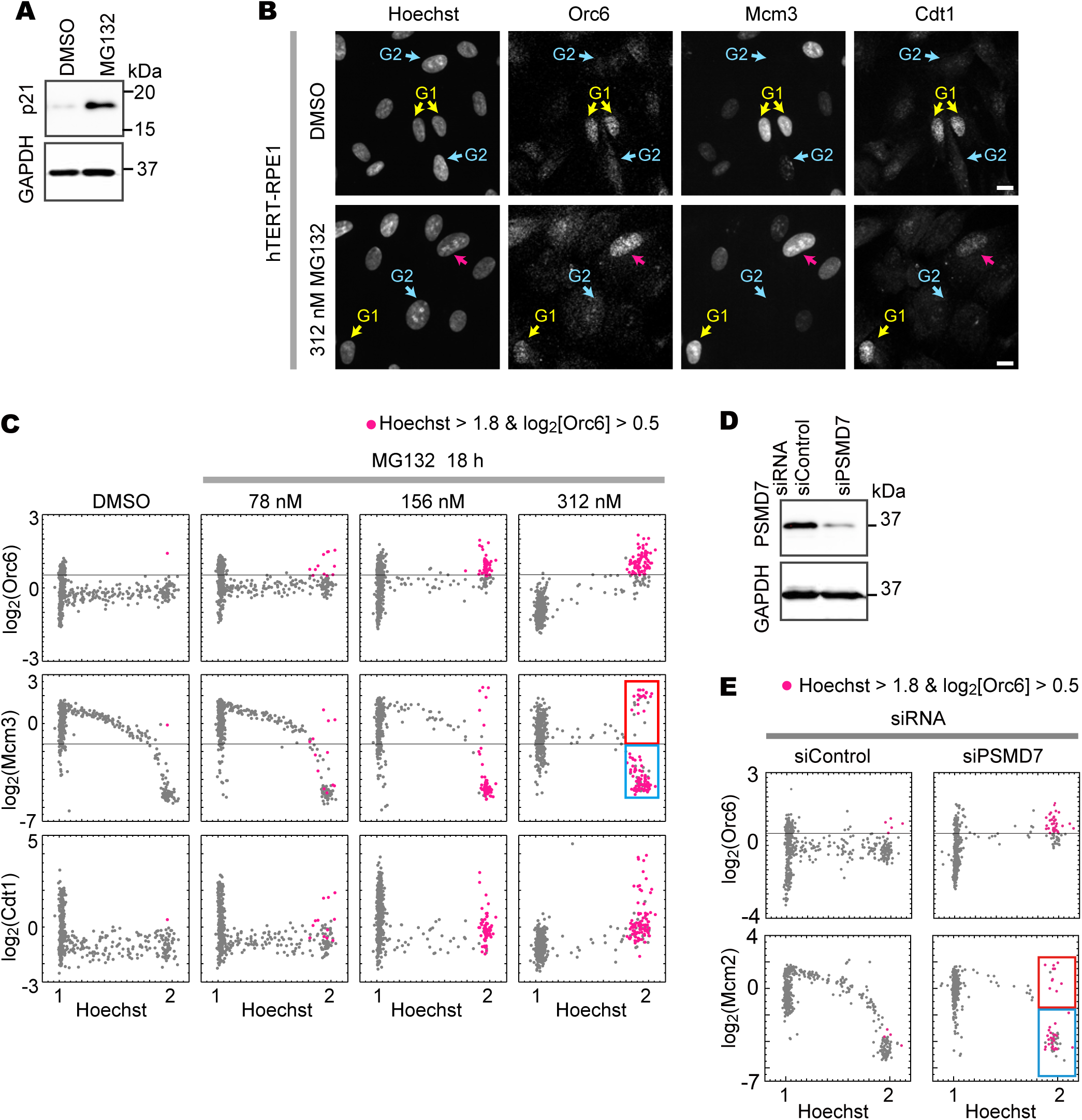
MG132 treatment increases the cells with high chromatin-bound Orc6 after the S-phase. (**A**) Western blot analysis of whole-cell lysates using antibodies against p21 (cell cycle marker) and GAPDH (loading control). hTERT-RPE1 cells were treated with DMSO or 312 nM MG132 for 1 day before harvesting. Molecular weight markers (kDa) are indicated on the right. (**B**) Representative fluorescence images of Hoechst, Orc6, Mcm3, and Cdt1 staining in hTERT-RPE1 cells treated with DMSO or 312 nM MG132 for 18 h. Cells were fixed using the pre-fixation method and stained with Hoechst, anti-Orc6, anti-Mcm3, and anti-Cdt1 antibodies. G1 (yellow arrows) and G2 (blue arrows) were labeled based on single-cell plot analysis in (**C**). Red arrows indicate a representative cell categorized in the red-boxed region in (**C**). Scale bars, 10 µm. (**C**) Single-cell plot analysis based on images of hTERT-RPE1 cells treated with DMSO or the indicated concentration of MG132, fixed and stained as described in (**B**). The cells with high Orc6 levels (log_2_[Orc6] > 0.5) and high DNA content (Hoechst > 1.8) are marked magenta in the Orc6 panels. The same cells are also marked magenta in the Mcm3 and Cdt1 panels. The red-boxed region represents cells with high Mcm3 levels and high DNA content, and the blue-boxed region represents cells with low Mcm3 levels and high DNA content. n = 450 (cells examined in each panel). (**D**) Efficiency of siRNA knockdown of PSMD7. Western blot analysis of hTERT-RPE1 cell lysates was performed with antibodies against PSMD7 and GAPDH (loading control) after 2 days of siRNA treatment. Molecular weight markers (kDa) are indicated on the right. (**E**) Single-cell plot analysis of Orc6 and Mcm2 in hTERT-RPE1 cells treated with siControl or siPSMD7. Cells were treated with siRNA for 2 days, fixed, and stained with Hoechst, anti-Orc6, and anti-Mcm2 antibodies. Cells with high Orc6 levels (log_2_[Orc6] > 0.5) and high DNA content (Hoechst > 1.8) are marked magenta in the Orc6 panels. The same cells are also marked magenta in the Mcm2 panels. The red-boxed region represents cells with high Mcm2 levels (log_2_[Mcm2] > ­1.5) and high DNA content (Hoechst > 1.8), and the blue-boxed region represents cells with low Mcm2 levels (log_2_[Mcm2] < ­1.5) and high DNA content (Hoechst > 1.8). n = 500 (number of cells examined in each panel). Another experiment with a different siRNA is shown in Fig. S4H.

Next, we examined the effect of low MG132 concentrations on Orc6 dynamics throughout the cell cycle. hTERT-RPE1 cells were treated with 78, 156, and 312 nM MG132 for 18 h. Cells were fixed using the pre-extraction method and stained with anti-Orc6, anti-Mcm3, and anti-Cdt1 antibodies (Fig. 4B), followed by single-cell plot analysis (Fig. 4C). We used anti-Mcm3 instead of anti-Mcm2 due to compatibility with immunostaining; previous single-cell plot analysis confirmed that chromatin-bound Mcm2 and Mcm3 exhibit similar cell cycle dynamics (Fig. S4F) (Hayashi-Takanaka et al., 2021). MG132 treatment produced cells with high levels of chromatin-bound Orc6 and high DNA content, marked as magenta dots in Orc6 panels (Orc6 >0.5 on the log_2_ scale, Hoechst >1.8, Fig. 4C). The number of these magenta cells increased with increasing MG132 concentrations, whereas few were observed in the control DMSO cells (Fig. 4C). These cells marked in Orc6 panels were also magenta-marked in the Mcm3 and Cdt1 panels (middle and lower panels, Fig. 4C). Some of these cells exhibited low Mcm3 signals (blue arrows in Fig. 4B; the blue-boxed region in Fig. 4C), while others exhibited high Mcm3 levels (red arrows in lower panels in Fig. 4B; red-boxed region in Fig. 4C). Co-staining with Cdt1 revealed that most magenta-marked cells also had relatively high levels of Cdt1 (Fig. 4C). Thus, the reduction of proteasome activity in hTERT-RPE1 cells appears to have led to the emergence of cells with increased loading of Orc6 and MCM on chromatin after replication, similar to the effect observed in passage 32 in IMR90 cells (Fig. 3).

As Cdt1 is a direct target of proteasome degradation, it is possible that chromatin loading of Orc6 and MCM depends on increased levels of Cdt1 under conditions of low proteasome activity. To examine this possibility, we inhibited Cdt1 degradation by treating cells with MLN4924, which inhibits the NEDD8-activating enzyme and thereby suppresses the function of cullin-RING E3 ubiquitin ligases (Soucy et al., 2009). Since Cdt1 degradation depends on these ligases, MLN4924 prevents its degradation (Lin et al., 2010). Treatment with 250 nM MLN4924 increased chromatin-bound Cdt1 levels during S-phase compared with the control (DMSO); however, it did not increase the chromatin-bound levels of Orc6 and MCM (Fig. S4G), suggesting that abnormal loading of Orc6 and MCM is not an indirect consequence of elevated Cdt1 levels.

To confirm the direct role of proteasomes in abnormal pre-RC formation after S-phase, we examined the effects of depleting PSMD7 — a subunit of the 26S proteasome essential for proteolytic activity (Shi et al., 2018) — using siRNA treatment. Cells were treated with PSMD7-targeting siRNA for 2 days, and Western blotting showed that PSMD7 levels were reduced to approximately 20% of those in control siRNA-treated cells (Fig. 4D). Under these conditions, cells were fixed using the pre-extraction method, stained with anti-Orc6 and anti-Mcm2 antibodies, and subjected to single-cell plot analysis (Fig. 4E, Fig. S4H). The results showed that PSMD7 depletion led to cells with high chromatin-bound Orc6 levels and high DNA content (magenta dots, Orc6 >0.5 on the log_2_ scale, Hoechst >1.8, Fig. 4E). Some of these cells had low Mcm2 levels (blue-boxed region, Fig. 4E), whereas others had high Mcm2 levels (red-boxed region, Fig. 4E), suggesting that proteasomal defects contribute to the abnormal loading of Orc6 onto chromatin after the S-phase. Taken together, these findings indicate that reduced proteasome activity promotes the reloading of Orc6 and MCM onto chromatin after S-phase.

### Inhibition of proteasome activity causes pre-RC formation without mitosis

To examine when cells with high levels of Orc6 and MCM, accompanied by high DNA content (red box in Fig. 4C), emerge following MG132 treatment, we performed a time-course analysis. We monitored cell cycle progression using anti-Mcm2 antibody and anti-histone H3 serine 10 phosphorylation (H3S10ph; an M-phase marker) antibody (Hayashi-Takanaka et al., 2009). In the case of control treated with DMSO, cells with high histone H3S10ph signals were observed (Fig. 5A; magenta dots, bottom left in Fig. 5B), indicating normal cell cycle progression. The percentage of histone H3S10ph-positive cells (magenta dots, ∼5% of total) is consistent with the duration of mitosis in hTERT-RPE1 cells from our previous study (∼1 h) (Hayashi-Takanaka et al., 2009; Hayashi-Takanaka et al., 2021). In contrast, after MG132 treatment, cells with high histone H3S10ph signals gradually disappeared by 8 h. During the first 4–12 h of treatment, cells with high DNA content (Hoechst ∼2) and low Mcm2 levels gradually increased (blue-circled regions, Fig. 5B), suggesting accumulation of G2-phase cells. Prolonged MG132 treatments (16–24 h) led to an increase in cells with high DNA content (Hoechst ∼2) and high Mcm2 levels (red-boxed regions, Fig. 5B), while no M-phase cells were detected. Using the same conditions, we analyzed the dynamics of chromatin-bound Orc6 and Cdt1. Chromatin-bound Orc6 levels increased in S-phase cells by 4 h after MG132 addition (upper panel in Fig. 5C), similar to the results observed with higher concentrations of MG132 (20 μM, Fig. 2). Chromatin-bound Cdt1 levels showed little change 4 h after treatment (Fig. 5C) but increased by 8 h in cells with Hoechst values around 2. The number of cells with high Orc6 and Cdt1 levels continued to increase by 12 h (orange-boxed regions, Fig. 5C), while cells with high MCM levels appeared later between 16 and 20 hours (red-boxed regions, Fig. 5B). We also performed experiments combining Mcm2 and Orc6 staining, and confirmed the timing of their chromatin loading (Fig. S6). These results suggest that Orc6 and Cdt1 begin to bind to chromatin within 8-12 hours after MG132 treatment, whereas MCM loading occurs later, between 16 and 24 hours after MG132 treatment.

**Figure 5.**
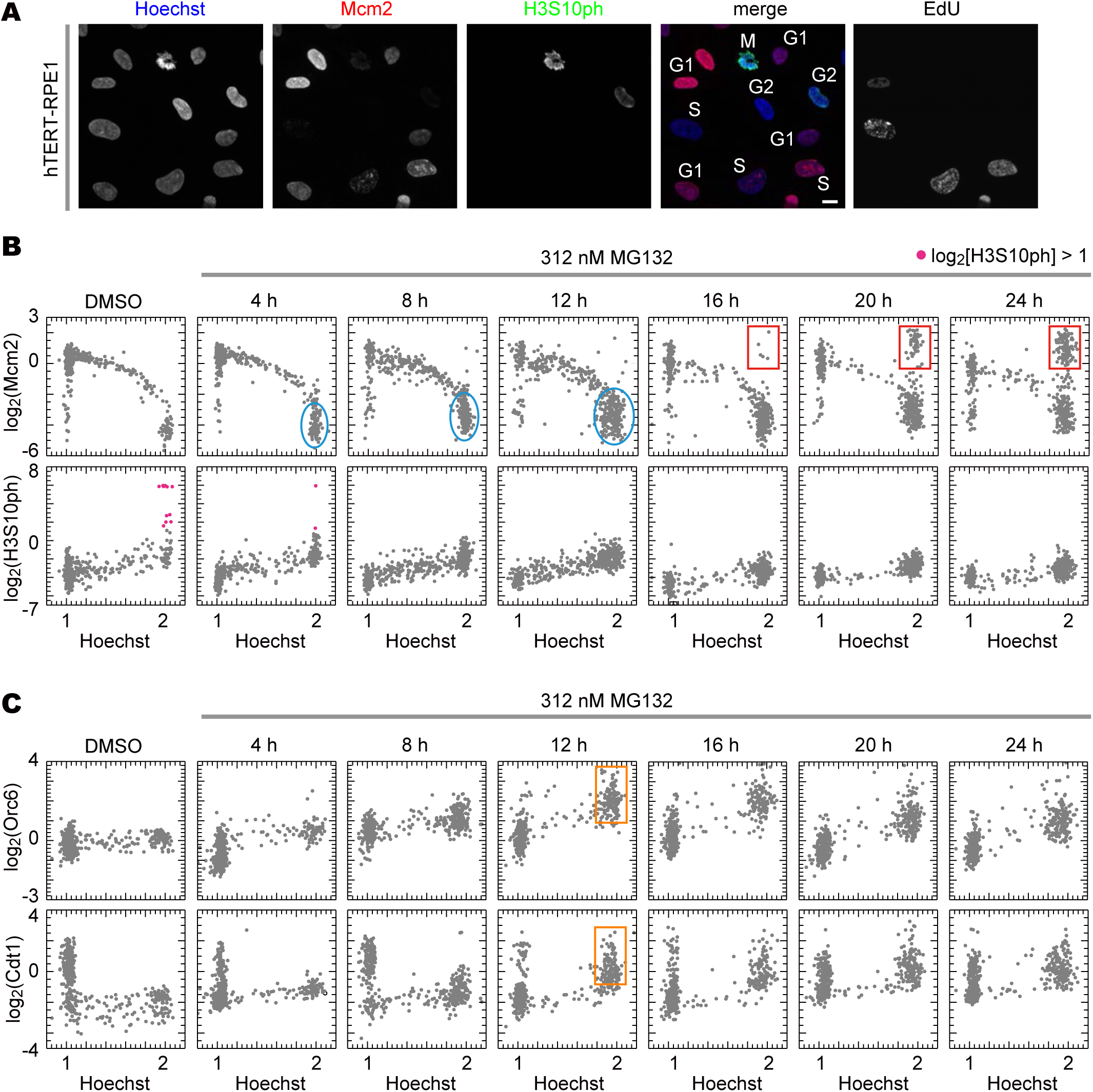
Time course of MG132-induced changes in Orc6, Mcm2 and Cdt1 indicate abnormal pre-RC formation without mitosis. (**A**) Images of Hoechst, Mcm2, histone H3S10ph, and EdU in the DMSO-treated cells fixed using the pre-extraction method. A combination of Mcm2, EdU and histone H3S10ph signals identifies cells in G1, S, G2, and M phases. The merged image shows Hoechst (blue), Mcm2 (red), and histone H3S10ph (green). Scale bar, 10 µm. (**B**) Single-cell plot analysis of Mcm2 (top), histone H3S10ph (bottom) in cells treated with DMSO or 312 nM MG132 for the indicated durations. Cells with high H3S10ph levels (log_2_[H3S10ph] > 1) are marked magenta in the H3S10ph panels. Blue-circled regions represent cells with low Mcm2 levels and high DNA content. Red-boxed regions represent cells with high Mcm2 and high DNA content. n = 425 (number of cells examined in each panel). (**C**) Single-cell plot analysis of Orc6 (top) and Cdt1 (bottom) in hTERT-RPE1 cells treated with DMSO or 312 nM MG132 for the indicated durations. Orange-boxed regions represent cells with high Orc6, Cdt1, and high DNA content in the 12-h treatment.

Taken together, we conclude that the cells with high DNA content (Hoechst ∼2) and elevated levels of Orc6 and Mcm2 represent tetraploid G1 cells that have arisen by skipping the M phase.

### Tetraploid G1 cells advance the cell cycle after release from proteasome inhibition

Next, to test whether these tetraploid G1 cells subsequently progress through the cell cycle, we monitored the dynamics of Orc6 and MCM during cell cycle progression following release from proteasome inhibition. In these experiments, cells were treated with 312 nM MG132 for 24 h (day 1), washed, and then cultured in MG132-free medium for an additional 24 h (day 2) (Fig. 6A). EdU was added to the culture medium for 30 min at the end of both days to label cells undergoing DNA replication. After 30-min labeling for each day, cells were fixed using the pre-extraction method and stained with anti-Mcm2, and anti-Orc6 antibodies (Fig. 6B), and subjected to single-cell plot analysis (Fig. 6C). EdU-positive cells were marked as orange dots, and the same cells were also identified in the Mcm2 and Orc6 panels (Fig. 6C). On day 1, the number of EdU-positive cells was limited, with most cells arrested at Hoechst levels of 1 or 2. Among the cells with Hoechst levels around 2 (Hoechst >1.8), a subset exhibited high levels of Mcm2 (Mcm2 >­1.5 on the log_2_ scale, red-boxed regions in Fig. 6C), accounting for 33.3 ± 3.7% of the total (mean and standard error of the mean in three independent experiments). On day 2, after MG132 removal, EdU-positive cells appeared in the Hoechst range of 2–4 (orange arrows in Fig. 6B; orange dots in day 2 panels in Fig. 6C), comprising 7.68 ± 0.40 % of the total (mean and standard error of the mean in three independent experiments). The Mcm2 pattern in cells with Hoechst levels 2–4 was similar to that in cells with Hoechst levels 1–2 (Mcm2 panel on day 2 in Fig. 6C), with Mcm2 levels increasing in G1-phase and decreasing during S-phase, reaching their lowest levels in G2-phase. These results suggest that cells with Hoechst levels of 2–4 (EdU panel on day 2 in Fig. 6C) underwent another round of DNA replication (i.e., whole-genome duplication) after MG132 removal. In the same single-cell plot in Fig. 6C, cells with high Orc6 levels (Orc6 > 0.5 on the log_2_ scale) and Hoechst values around 2 were marked as magenta dots in the Orc6 panels, and the same cells were also marked as magenta dots in the Mcm2 panels (Fig. 6C’). As observed in Fig. 4D, on day 1, the population of cells marked as magenta dots (high Orc6 and Hoechst ∼2) split into two groups (Fig. 6C’): one with high Mcm2 levels and the other with relatively low Mcm2 levels (red and blue arrows in Fig. 6B, respectively; red- and blue-boxed regions in Fig. 6C’, respectively). On day 2, however, most of the magenta-marked cells exhibited high Mcm2 levels, being consistent with the result that Orc6 is loaded onto chromatin prior to Mcm2, as shown in Fig. 5B, C. Taken together, these results confirm that, after release from proteasome inhibition, cells with high MCM and Orc6 levels skip mitosis and proceed with an additional round of DNA replication to continue the tetraploid cell cycle.

**Figure 6.**
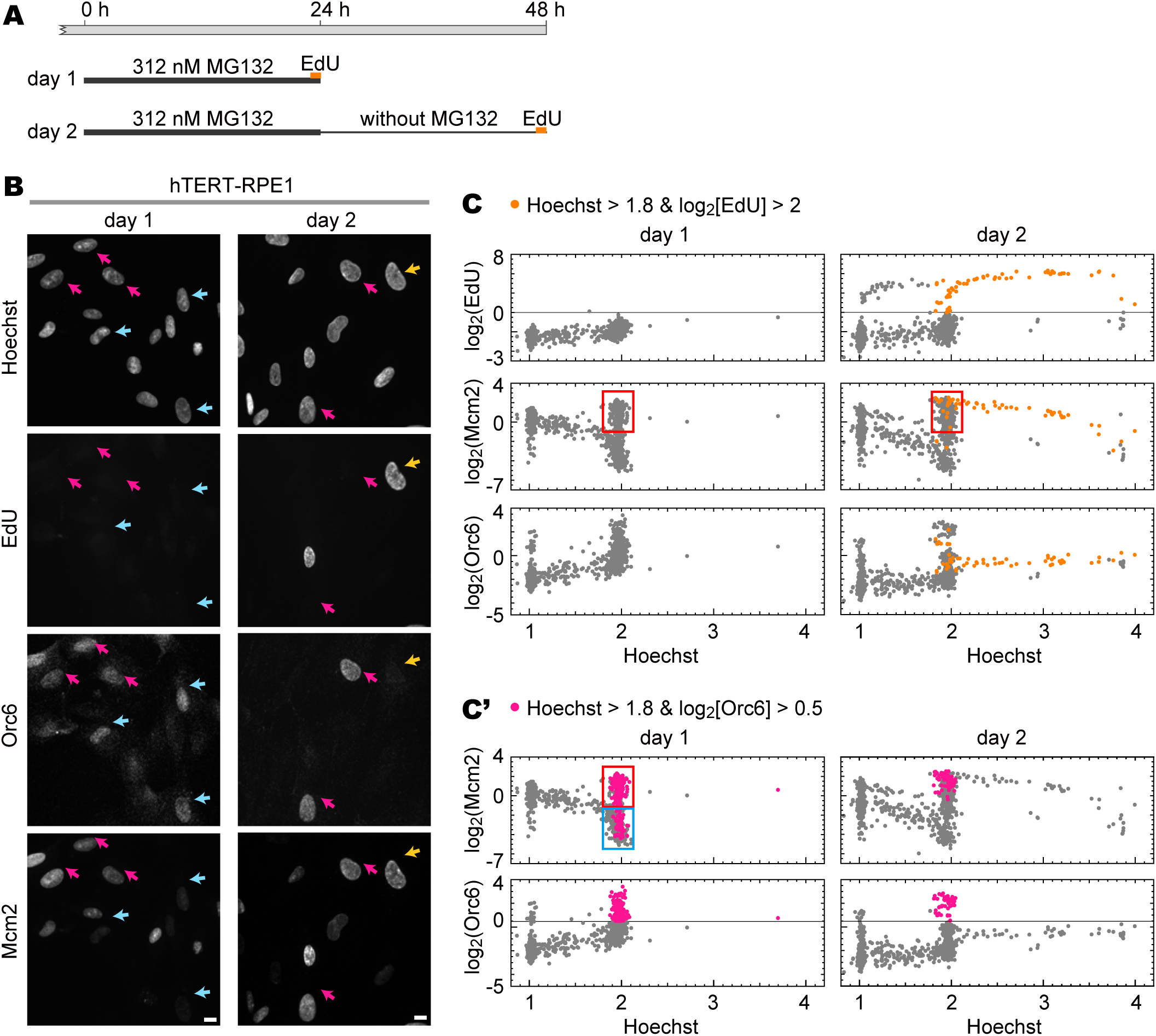
Cells forming pre-RC via proteasome inhibition can progress through the cell cycle as tetraploids. (**A**) Timing diagram for MG132 treatment and EdU labeling used in (**B)** and **(C**). hTERT-RPE1 cells were incubated with 312 nM MG132 for a day (day 1) (left panels in **B**), washed with fresh medium without MG132, and incubated for another day (day 2) (right panels in **B**). The cells were labeled with EdU for 30 min before the sample collection on day 1 and day 2. (**B**) Representative fluorescence images of Hoechst, Orc6, and Mcm2 staining used in single-cell plot analysis in (**C** and **C’**). Red and blue arrows indicate typical cells in the red and blue boxes in (**C** and **C’**). Orange arrows indicate a tetraploid S-phase cell (orange dots of day 2 in **(C)**). Scale bars, 10 µm. (**C**) Single-cell plot analysis of EdU (top), Mcm2 (middle), and Orc6 (bottom). The orange dots represent S-phase cells based on EdU intensities (log_2_[EdU] > 2) with high DNA contents (Hoechst > 1.8). The red-boxed regions represent cells with high Mcm2 levels (log_2_[Mcm2] > ­1.5) and high DNA contents (1.8 < Hoechst < 2.1). n = 600 (cells examined in each panel). (**C’**) The same data from (**C**) is reused but marked differently. Cells with high Orc6 levels (log_2_[Orc6] > 0.5) and high DNA content (Hoechst > 1.8) are marked magenta in the Orc6 panels. The same cells are also marked magenta in the Mcm2 panels. The red-boxed region represents cells with high Mcm2 levels (log_2_[Mcm2] > ­1.5) and high DNA content (1.8 < Hoechst < 2.1). The blue-boxed region represents cells with low Mcm2 levels (log_2_[Mcm2] < -1.5) and high DNA content (1.8 < Hoechst < 2.1).

### Involvement of Orc6 in MCM loading in mitosis-skipped cells

To determine whether the presence of Orc6 on chromatin after S-phase entry contributes to MCM loading, we examined chromatin-bound MCM levels following Orc6 depletion by siRNA in hTERT-RPE1 cells (Fig. 7A). As a control, we also performed siRNA depletion of Orc1 (Fig. S6), a subunit essential for MCM loading (Coulombe et al., 2019; Ogawa et al., 1999; Samson et al., 2016) that is known to be degraded in a proteasome-dependent manner after S-phase entry (Li and DePamphilis, 2002; Méndez et al., 2002; Tatsumi et al., 2003). Western blot analysis and single-cell plot analysis confirmed the efficient depletion of Orc6 in whole-cell and chromatin-bound fractions after 42 h siRNA treatment (Fig. 7B, Fig. S6). Under these conditions, siRNA-treated cells were fixed using the pre-extraction method, stained with Hoechst and anti-Mcm2 antibody, and subjected to single-cell plot analysis. Single-cell plot analysis showed that cells treated with siOrc1 or siOrc6 accumulated at G1 phase (81.0% and 71.3%, middle and right panels, respectively in Fig. 7C) compared with control siRNA (66.2%, left panel in Fig. 7C). To quantify MCM loading, cells with high MCM levels (cyan dots, half of the maximum value of Mcm2 in Hoechst <1.125, Fig. 7C) were expressed as a percentage of the cells with Hoechst signals >1.125 (Fig. 7D). The percentage of cells with high MCM levels was significantly reduced in Orc6-depleted cells (9.6 ± 2.3%) and in Orc1-depleted cells (3.8 ± 0.8%) compared to in siControl (26.4 ± 6.8%) (Fig. 7D). These results suggest that Orc6 plays a role in MCM loading in the normal G1 phase although its effect is less pronounced than that of Orc1.

**Figure 7.**
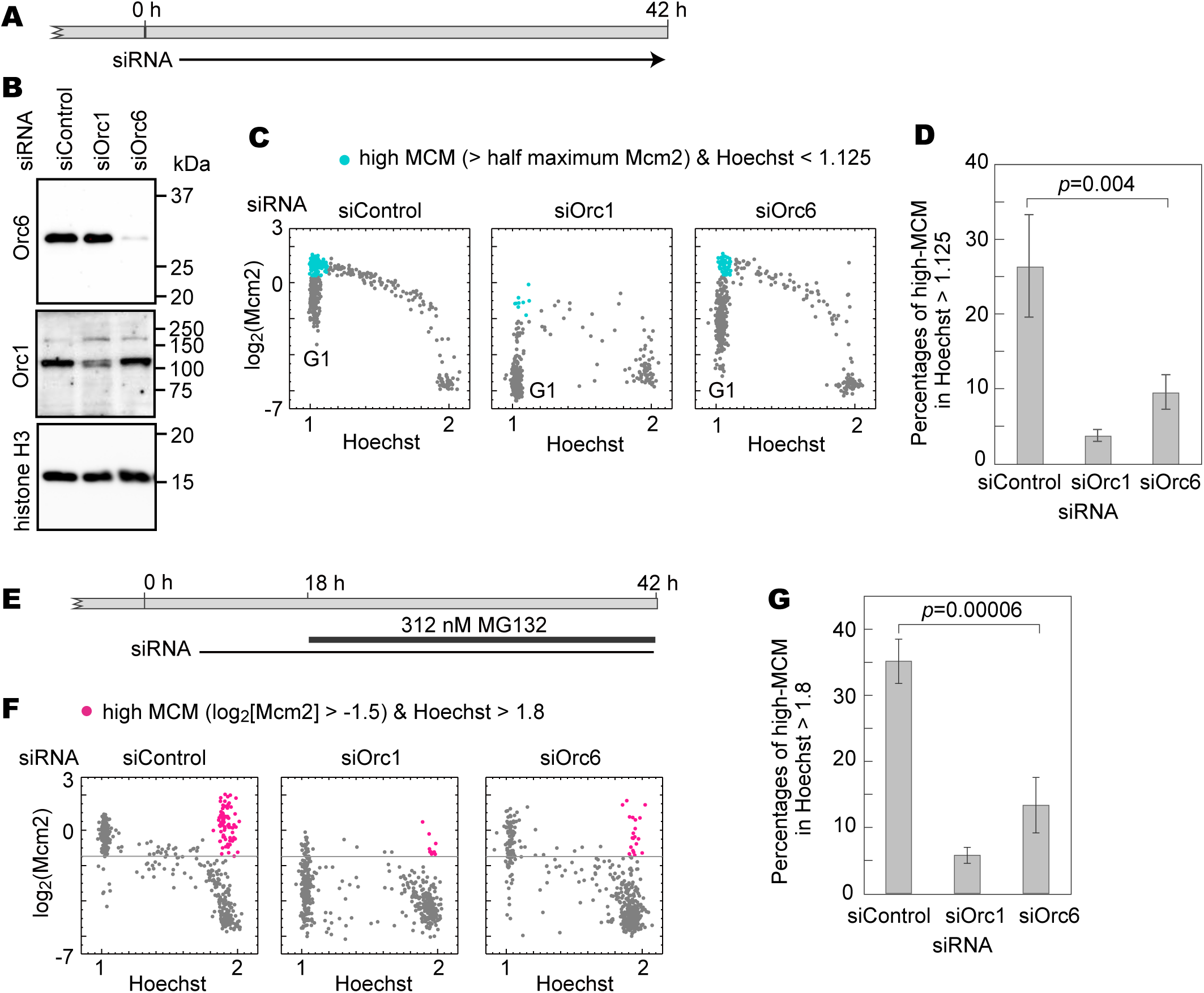
Effects of Orc6 depletion on the formation of tetraploid cells. (**A**) Schematic diagram of siRNA treatment timing. hTERT-RPE1 cells were treated with siRNA targeting Orc6 (siOrc6) for 42 h. Conditions for Orc6 siRNA was determined as shown in Fig. S6C. (**B**) Western blot analysis of whole-cell fractions from cells treated with siRNA against Orc6 (siOrc6), together with a negative control (siControl), and Orc1 as a possitive control (siOrc1), using antibodies against Orc6, Orc1, and histone H3. Molecular weight markers (kDa) are indicated on the right. (**C**) Single-cell plot analysis of Mcm2 in hTERT-RPE1 cells treated with siRNA (siControl, siOrc1, or siOrc6). siRNA-treated cells were fixed using the pre-extraction method and stained with Hoechst and anti-Mcm2 antibody. Cells with high Mcm2 levels (> half the maximum Mcm2 value, excluding the 4 highest values among 450 cells) and low DNA content (Hoechst < 1.125) are marked cyan. n = 450 (number of cells examined in each panel). **(D)** Percentages of cyan-marked cells (high MCM) among G1-phase cells (Hoechst < 1.125) based on (**C**). Columns and error bars represent the mean and standard error of the mean of three independent experiments. Statistical difference (*p* = 0.004) was determined using a one-sided *t-*test. (**E**) Timing diagram for siRNA treatment and MG132 addition. hTERT-RPE1 cells were treated with siRNA. After 18 h of siRNA treatment, MG132 was added at 312 nM for an additional 24 h, with continued siRNA treatment. (**F**) Single-cell plot analysis of Mcm2 in hTERT-RPE1 cells treated with siRNA (siControl, siOrc1, or siOrc6), with MG132 addition. Cells with high Mcm2 levels (log_2_[Mcm2] > ­1.5) and high DNA content (Hoechst > 1.8) are marked magenta. n = 450 (number of cells examined in each panel). Another experiment with different siRNA is shown in Fig. S7A. (**G**) Percentage of magenta-marked cells (log_2_[Mcm2] > ­1.5 and Hoechst > 1.8) among G2-phase cells (Hoechst >1.8) based on (**F**). Columns and error bars represent the mean and standard error of the mean of four independent experimental results. Statistical differences (*p* = 0.0006) were determined using a one-sided *t-*test.

We next examined the effect of Orc6 depletion on Mcm2 loading during the tetraploid G1 phase induced by proteasome inhibition. Cells were treated with siRNA for 18 h, followed by MG132 treatment for 24 h while maintaining siRNA treatment (Fig. 7E). After treatment, cells were fixed using the pre-extraction method, stained with Hoechst and anti-Mcm2 antibody, and analyzed using single-cell plot analysis (Fig. 7F; Fig. S7A). Cells with doubled DNA content (Hoechst ∼2) accumulated by MG132 treatment in all siControl, siOrc1 and siOrc6 (Fig. 7F). Cells with high MCM levels and Hoechst ∼2 (magenta dots, Hoechst >1.8 and Mcm2 >­1.5 on the log_2_ scale, Fig. 7F) were quantified as a percentage of the total cells with Hoechst >1.8 (Fig. 7G). Orc6 depletion significantly reduced the percentage of high-MCM cells to 13.3 ± 4.2% compared to the siControl (35.2 ± 3.4%) (Fig. 7G), underscoring the importance of Orc6 in MCM loading in tetraploid G1 cells, while Orc1 depletion reduced the percentage of high-MCM cells (5.7 ± 1.2%). We also examined a role of Cdc6, another pre-RC component, in MCM loading after siRNA treatment. Single-cell plot analysis revealed a reduction in tetraploid G1 cells with high MCM levels (magenta dots, Hoechst >1.8, and Mcm2 >−1.5 on the log_2_ scale) after Cdc6 depletion (Fig. S7B, C, D) to a similar level to Orc6 depletion. These results indicate that the presence of Orc6, as well as Cdc6, on chromatin contributes to MCM reloading in tetraploid G1 cells, although their influence is less prominent than that of Orc1.

In summary, during G1-phase, Orc6 binds to chromatin along with other pre-RC components. Upon S-phase entry, Cdt1 and Orc1 are degraded by proteasome activity, while Orc6 dissociates from chromatin (Fig. 8A). Orc2–5 remain bound to chromatin throughout the cell cycle. These cells proceed through mitosis, producing normal diploid cells (Fig. 8A). However, in cells with reduced proteasome activity, Orc6, Orc1, Cdt1, and MCM are reloaded onto chromatin. These cells bypass mitosis and proceed into replication, leading to tetraploid formation (Fig. 8B).

**Figure 8.**
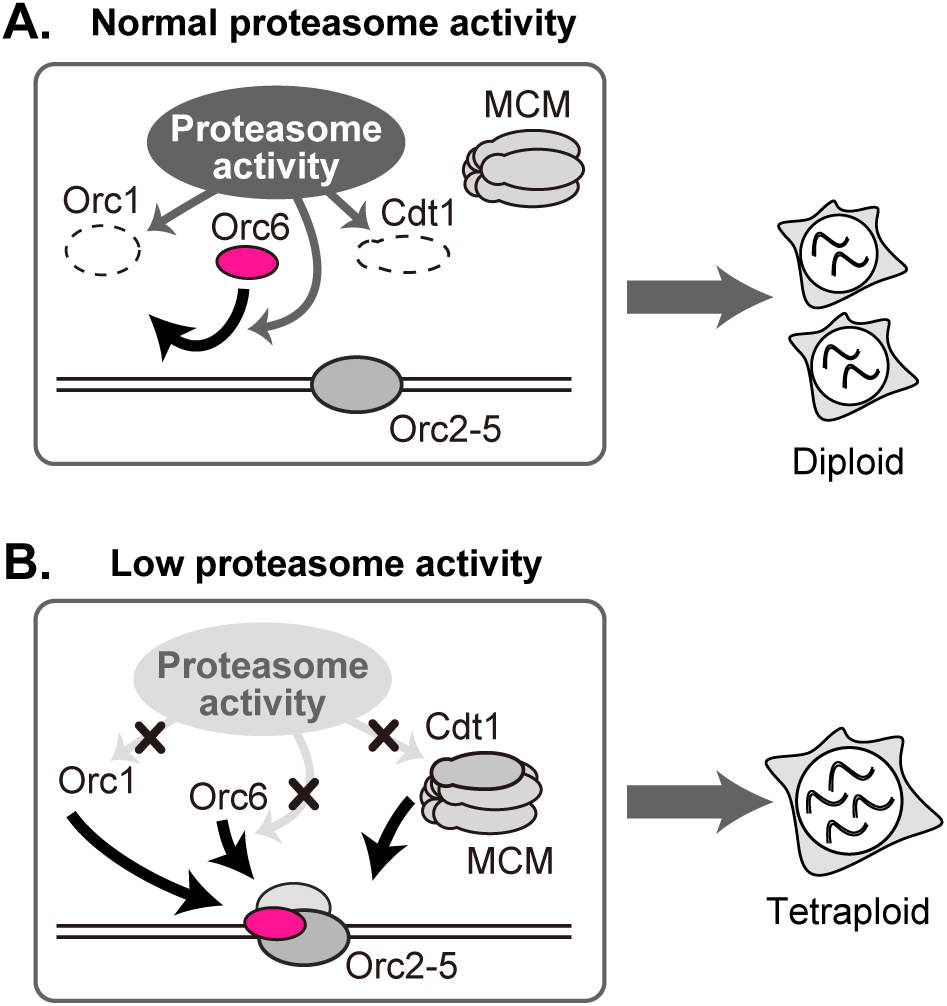
Schematic diagram of Orc6 behavior. **(A)** Normal proteasome activity: After DNA replication, Cdt1 and Orc1 are degraded by proteasome activity, and Orc6 dissociates from chromatin, preventing MCM reloading. Cells enter mitosis and produce normal diploid cells. Orc2–5 represents a complex of Orc2, Orc3, Orc4, and Orc5. (**B**) Loss of proteasome activity: Orc6, Orc1, Cdt1, and MCM are reloaded onto chromatin without mitosis, allowing cells to re-enter the replication phase as tetraploid cells.

## Discussion

In this study, we demonstrated dynamic changes in chromatin-bound Orc6 levels during the cell cycle in hTERT-RPE1 cells. The levels of chromatin-bound Orc6 increased until the G1/S transition, decreased upon entry into the S-phase, and remained low until the next G1-phase, despite the total amount of Orc6 remaining almost unchanged throughout the cell cycle. Contrary to our results, previous western blot analysis using synchronized HeLa cells showed that chromatin-bound Orc6 levels remained nearly constant across different cell cycle phases (Méndez et al., 2002). However, our single-cell plot analysis revealed a more pronounced difference in Orc6 levels between G1 and S/G2 phases in normal cell lines (IMR90, and hTERT-RPE1 cells) compared to cancer cell lines (HeLa and U2OS cells) (Fig. 1, Fig. S2). This suggests that the regulation of chromatin-bound Orc6 during the cell cycle could serve as a distinguishing factor between normal and cancerous cells.

We further demonstrated that the removal of Orc6 from chromatin after the S-phase, through a proteasome-dependent mechanism, plays a role in preventing abnormal MCM loading. Aberrant MCM loading beyond the S-phase can initiate an additional round of DNA replication without mitotic division, leading to tetraploidy, which may contribute to cancer development. To prevent such occurrences, the ubiquitin-proteasome system regulates the levels of several pre-RC factors, such as Cdt1, Cdc6, and Orc1, at specific cell cycle phases (Li and DePamphilis, 2002; Méndez et al., 2002; Nishitani et al., 2006; Tatsumi et al., 2003; Walter et al., 2016). Interestingly, unlike other pre-RC factors, Orc6 is not degraded at the protein level but is instead dissociated from chromatin via proteasome activity. This indicates that the proteasome regulates pre-RC protein levels on chromatin in a distinct manner, thereby preventing abnormal MCM loading. Certain carcinomas exhibit elevated Orc6 levels (Pan et al., 2022; Tang et al., 2023), suggesting that excess Orc6 might interfere with its chromatin dissociation, potentially leading to the reloading of pre-RC onto chromatin. Our results also suggest that eliminating Orc6 from chromatin at the onset of the S-phase is critical. Given that Orc6 is the most evolutionarily diverse ORC subunit (Dhar and Dutta, 2000), its dissociation from chromatin during the S-phase may represent an auxiliary mechanism uniquely evolved to prevent aberrant MCM loading.

The precise molecular mechanism driving Orc6 dissociation from chromatin remains unclear. Orc6 lacks known ubiquitinated amino acid residues for subsequent proteasome degradation (Coulombe et al., 2019). One possibility is that the chromatin-bound Orc6 levels are regulated by factors degraded through proteasome activity, such as Orc1 or Cdt1. However, our experiments showed that reducing Orc1 and Cdt1 levels did not affect the chromatin-bound Orc6 levels. Another plausible explanation involves post-translational modifications, such as phosphorylation, as observed with *S. cerevisiae* Orc6 (Chen and Bell, 2011; Chen et al., 2007). Notably, S-phase-specific phosphorylation of Orc6 has been shown to contribute to genome stability via DNA damage checkpoints (Lin et al., 2023). These modifications may facilitate Orc6 dissociation from chromatin, possibly mediated by the degradation of modifying enzymes through proteasome activity. In addition, proteasomes may have non-proteolytic functions. For example, the ATPase activity within the proteasome can unwind DNA (Makino et al., 1996; Makino et al., 1999), suggesting that a proteasome lacking proteolytic activity might still facilitate chromatin binding by ORC, including Orc6. Furthermore, inhibition of proteasome activity has been associated with nucleosome hyperacetylation at transcription start sites (Kinyamu et al., 2020), which overlaps with replication origins (Sugimoto et al., 2018). This hyperacetylation may promote Orc6 binding to chromatin.

A decline in proteasome activity affects a variety of cellular processes, including senescence (Huang et al., 2022; Vilchez et al., 2014). Recent studies show that cellular senescence is associated with a tetraploid state in G1-phase, which occurs when cells bypass M-phase (Johmura et al., 2014; Panopoulos et al., 2014; Zeng et al., 2023). In agreement, we observed that proteasome inhibition by MG132 led to tetraploid G1 cells, and subsequent removal of the inhibitor restored proteasome activity, promoting cell progression into S-phase (Figs. 4, 5). This implies that age-related declines in proteasome activity may contribute to the production of tetraploid G1 cells, increasing the risk of cancer development.

In conclusion, our results show that normal cells can progress to a tetraploid G1-phase when proteasome activity is reduced, a key step linked to whole-genome duplication and carcinogenesis (Bielski et al., 2018; Gemble et al., 2022; López et al., 2020). This study contributes to our understanding of the interplay between replication factor dynamics and proteasome activity, further connecting these processes to tetraploid cell formation, senescence, and cancer development.

## Materials and Methods

### Antibodies for immunofluorescence staining and Western blotting

The primary antibodies used were anti-Orc6 (1:300 for immunofluorescence staining (IF) and 1:1000 for Western blot (WB); RRID:AB_670295, 3A4, sc-32735; Santa Cruz Biotechnology), anti-Orc2 (1:300 for IF and 1:1000 for WB; RRID:AB_592395, M055-3, MBL), anti-Cdt1 (1:300 for IF and 1:1000 for WB; RRID:AB_10896851, D10F11; Cell Signaling Technology), anti-Mcm3 (1:1000 for IF; sc-390480; Lot#J1118, Santa Cruz Biotechnology; 1:300 (IF); RRID:AB_398024, Anti-BM28; 610701, BD Transduction), anti-Mcm2 (1:1000 for IF and WB; RRID:AB_2142137, D7G11, Cell Signaling Technology), anti-Orc1 (1:100 for IF and 1:300 for WB; F-10, sc-398734, Santa Cruz Biotechnology), anti-GAPDH (1:1000 for WB; RRID:AB_1903993, 14C10; Cell Signaling Technology), anti-histone H3 (1G1, 1:1000 for WB) (Nozawa et al., 2010), anti-histone H3S10ph (1:2000 for IF; CMA311, (Hayashi-Takanaka et al., 2009)), anti-PSMD7 (1:1000 for IF and WB; EPR13517, ab181072; Abcam), anti-Cdc6 (1:300 for IF and 1:1000 for WB,RRID:AB_627236, sc-9964, Santa Cruz Biotechnology), and anti-p21 (1:1000 for IF and WB; RRID:AB_823586, 12D1; Cell Signaling Technology). The specificity of these antibodies used for immunostaining is shown in Fig. S1. We previously confirmed by single-cell plot analysis based on immunostaining that chromatin-bound Mcm2 and Mcm3 showed similar dynamics during the cell cycle (Hayashi-Takanaka et al., 2021).

For fluorescent staining, the secondary antibodies were as follows: Alexa 488 donkey anti-mouse IgG (RRID:AB_2341099, 715-545-151; Jackson ImmunoResearch), Cy5 donkey anti-mouse IgG (RRID:AB_2340820, 715-175-151; Jackson ImmunoResearch), Cy3 donkey anti-Rat IgG (RRID:AB_2340667, 712-165-153; Jackson ImmunoResearch), Alexa 488 donkey anti-rabbit IgG (RRID:AB_2313584, 711-545-152; Jackson ImmunoResearch) and Cy3 donkey anti-rabbit IgG (RRID:AB_2340585, 711-005-152; Jackson ImmunoResearch).

### Cell culture and chemicals

hTERT-RPE1 (ATCC, CRL-4000), IMR90 (ATCC, CCL-186), U2OS (ATCC, HTB-96), and HeLa (RCB0007) cell lines were grown in a culture medium containing high-glucose Dulbecco’s modified Eagle’s medium (Sigma-Aldrich) supplemented with penicillin/streptomycin (100 units/mL penicillin, 100 μg/mL streptomycin; Fujifilm) and 10% fetal calf serum (Thermo Fisher Scientific), as previously described. Unless otherwise noted, the experiments were performed using hTERT-RPE1 cells.

siRNA transfection was performed using Lipofectamine RNAiMAX (Thermo Fisher Scientific) with siRNAs (SASI_Hs01_00069342 for Orc6, SASI_Hs01_00069343 for Orc6#2, SASI_Hs01_00038888 for Orc1, SASI_Hs02_00341067 for Orc2, and Universal Negative Control #1, Sigma-Aldrich; s11403 for PSMD7, s11401 for PSMD7#2, and s2745 for Cdc6, Ambion, Thermo Fisher Scientific) following the manufacturer’s protocol.

MG132 (Fujifilm) and MLN4924 (MedChemExpress) were dissolved in DMSO.

### Fluorescence staining for single-cell plot analysis

Immunostaining was performed as previously described (Hayashi-Takanaka et al., 2011; Hayashi-Takanaka et al., 2015; Hayashi-Takanaka et al., 2021). Briefly, the cells (∼2 × 10^4^ cells/cm^2^) were plated on a 12-well plate with a coverslip (15 mm diameter; No. 1S; Matsunami) at the bottom and cultured for over 1 day.

In the direct fixation method, cells were first fixed with 1 mL of the fixative (2% paraformaldehyde [Electron Microscopy Sciences] dissolved in 250 mM HEPES [pH 7.4]) for 10 min at room temperature (approximately 25°C), followed by permeabilization for 15 min using 1 % Triton X-100 in PBS, and subsequently blocked with Blocking One (Nacalai Tesque, Inc.).

In the pre-extraction method, before fixation, cells were first treated with 1 mL of 0.2% Triton X-100 solution containing 20 mM HEPES (pH 7.4), 100 mM NaCl, and 300 mM sucrose for 3 min on ice. After removing the Triton X-100 solution, the cells were fixed using the fixative described above. This fixation method removes the nuclear membranes and loosely bound chromatin proteins.

The fixed cells were incubated with specific primary antibodies (0.2‒1 μg/mL) for 2 h at room temperature (approximately 25°C) and washed three times (10 min each) using PBS. When cells were stained with multiple antibodies, the cells were simultaneously incubated with these antibodies. The cells were treated with secondary antibodies (1 μg/mL) and Hoechst (0.1 μg/mL) for 2 h at room temperature (approximately 25°C) and subsequently washed three times with PBS. The cells were mounted using the ProLong Glass Antifade Mountant (Thermo Fisher Scientific).

The replication foci were labeled by culturing the cells in a medium containing 10 μM 5-ethynyl-2′-deoxyuridine (EdU) for 30 min. Subsequently, the cells were fixed using the direct fixation or pre-extraction method, and the resulting EdU signals were detected with Alexa Fluor 647 using a Click-iT EdU Imaging Kit (Thermo Fisher Scientific).

For staining of SA-βGal (senescence-associated β-galactosidase), live cells were treated with the reagents according to commercial protocols (Cellular Senescence Detection Kit, SPiDER-βGal. Dojindo).

### Microscopy

Fluorescence images were obtained using a DeltaVision Elite system (GE Healthcare Inc.) equipped with a pco.edge 4.2 sCMOS camera (PCO) and an Olympus 40× UApo/340 oil immersion objective lens (NA 0.65 with iris) with DeltaVision Elite Filter Sets (DAPI-FITC-TRITC-Cy5).

### Single-cell plot analysis

Single-cell plot analysis was used to measure the levels of multiple intracellular components based on fluorescence microscopic images and plot their correlation in individual cells (Hayashi-Takanaka et al., 2020; Hayashi-Takanaka et al., 2021). The fluorescence intensity of each nucleus was measured using NIS Elements software (version 3.0; Nikon). The background intensity outside the cells was subtracted. The nuclear areas of individual cells were determined by automatic thresholding using Hoechst signals. The sum of the intensities (i.e., the average intensity × nuclear area) in each nucleus was measured across all the fluorescence channels. Cells in the M-phase were excluded from analysis except those shown in Fig. 5B, 5C. In Fig. 5B, 5C, M-phase cells were used for the analysis. To plot the Hoechst intensity distribution of 450– 600 nuclei, the signal intensity in each nucleus was normalized relative to those with the 20th lowest and 20th highest intensities, designated as 1 and 2, respectively, because nuclei with the lowest and highest intensities were sometimes outliers. Hoechst intensity was plotted on a linear scale. The average intensities of Orc6, EdU, Orc2, Cdt1, Mcm2, and Mcm3 were set to 1 (0 on the log_2_ scale) and plotted on the log_2_ scale. The fluorescence intensities were normalized to time “0 h” (Figs. 2A, 2B), DMSO treatment (Figs. 4C, 5B, 5C, S4G, S5), the negative control siRNA (Figs. 4E, 7C, 7F, S4H, S6A, S6B, S7A, S7C) and MG132 treatment for day 1 (Fig. 6C). The values relative to the average were plotted using Mathematica (version 13.0) (Wolfram Research).

### Preparation of chromatin-bound fractions and whole-cell lysates

Native chromatin-bound fractions (hereafter, referred to as “chromatin-bound fractions”) were prepared by the method previously described (Quan et al., 2015; Sheu and Stillman, 2006) with some modifications (Hayashi-Takanaka et al., 2021). The cells were washed with cold PBS, suspended in buffer A (50 mM HEPES [pH 7.4], 100 mM KCl, 2.5 mM MgCl_2_, 0.1 mM ZnSO_4_, 1 mM ATP, 1 mM dithiothreitol [DTT], and protease inhibitor cocktail [EDTA-free, Nacalai Tesque, Inc.]), and collected via centrifugation (1,300 *g*, 3 min, 4°C). The cells were subsequently treated with 200 µL of 0.2% Triton X-100 in buffer A, layered with 200 µL of 30% sucrose cushion in buffer A containing 0.2% Triton X-100, and centrifuged at 20,000 × *g* for 10 min at 4°C. The pellets were collected as chromatin-bound fractions and dissolved in 1× SDS loading buffer. The suspension was then subjected to SDS-PAGE.

To prepare whole-cell lysates, the cells were washed with PBS and suspended in an equal volume of 2× SDS loading buffer. The suspension was subjected to SDS-PAGE.

### Western blot analysis

Chromatin-bound fractions and whole-cell lysates were suspended in 1× SDS loading buffer and subjected to SDS-PAGE (10%, 12%, or 15%) to separate proteins. Proteins were transferred onto Immobilon-P PVDF membranes (Merck) using a semi-dry blotting system (ATTO). After blocking with 5% skim milk (Nacalai Tesque, Inc.), the membranes were incubated with the primary antibodies. After incubation with peroxidase-conjugated secondary antibodies (GE Healthcare), signals were detected using ImmunoStar LD (Fujifilm Wako) and imaged with the ChemiDoc imaging system (BioRad).

To quantify Orc6 protein levels, quantitative Western blotting was performed as shown in Fig. S6C. For cells treated with control siRNA, a fraction corresponding to 4 × 10^4^ cells was defined as 1 and serially diluted to 0.5, 0.25, 0.125, and 0.0625. After signal detection using the ChemiDoc image system (BioRad), band intensities were quantified with background subtraction using Image Lab software (BioRad). Linearity was confirmed within this dilution range (Fig. S6C). In cells treated with siORC6, Orc6 levels in the fraction corresponding to 4 × 10^4^ cells were approximately 0.2 in the whole-cell fraction and 0.4 in the chromatin-bound fraction, relative to the control (Fig. S6C).

To examine the effect of MG132 treatment on Orc6 protein levels, three independent experiments were conducted. Western blot analysis was performed for each experiment (left in Fig. S3C). Signals were detected using the ChemiDoc image system (BioRad) and quantified using Image Lab software (BioRad), as described above. Signal intensities were normalized to the band intensity of the untreated whole-cell fraction (0 h, leftmost), which was set to 1. Normalized intensities of each band were plotted (right in Fig. S3C) and the mean values from the three experiments are indicated by horizontal bars.

## Acknowledgements

We thank Ms. Chizuru Ohtsuki and Yoshino Kubota for their technical assistance, and Drs. Hisao Masai, Haruhiko Asakawa, and Yasuhiro Hirano for insightful discussion. We also thank Dr. Hiroshi Kimura for histone H3 antibodies and Dr. Hisao Masukata for critically reading the manuscript.

This work was supported by JSPS KAKENHI grants: JP18H05530, JP20K20582 and JP20K08698 (to IH); JP18H05533, JP20H00454 and JP23K05636 (to YH); and JP18H05528 (to TH). This work was also supported by the Naito Foundation, Urakami Foundation for Food and Food Culture Promotion, and the Foundation for Promotion of Cancer Research in Japan (to YHT).

## Data Availability

Source data are available from the corresponding authors upon reasonable request.

## Supplemental Figures

**Figure S1.**
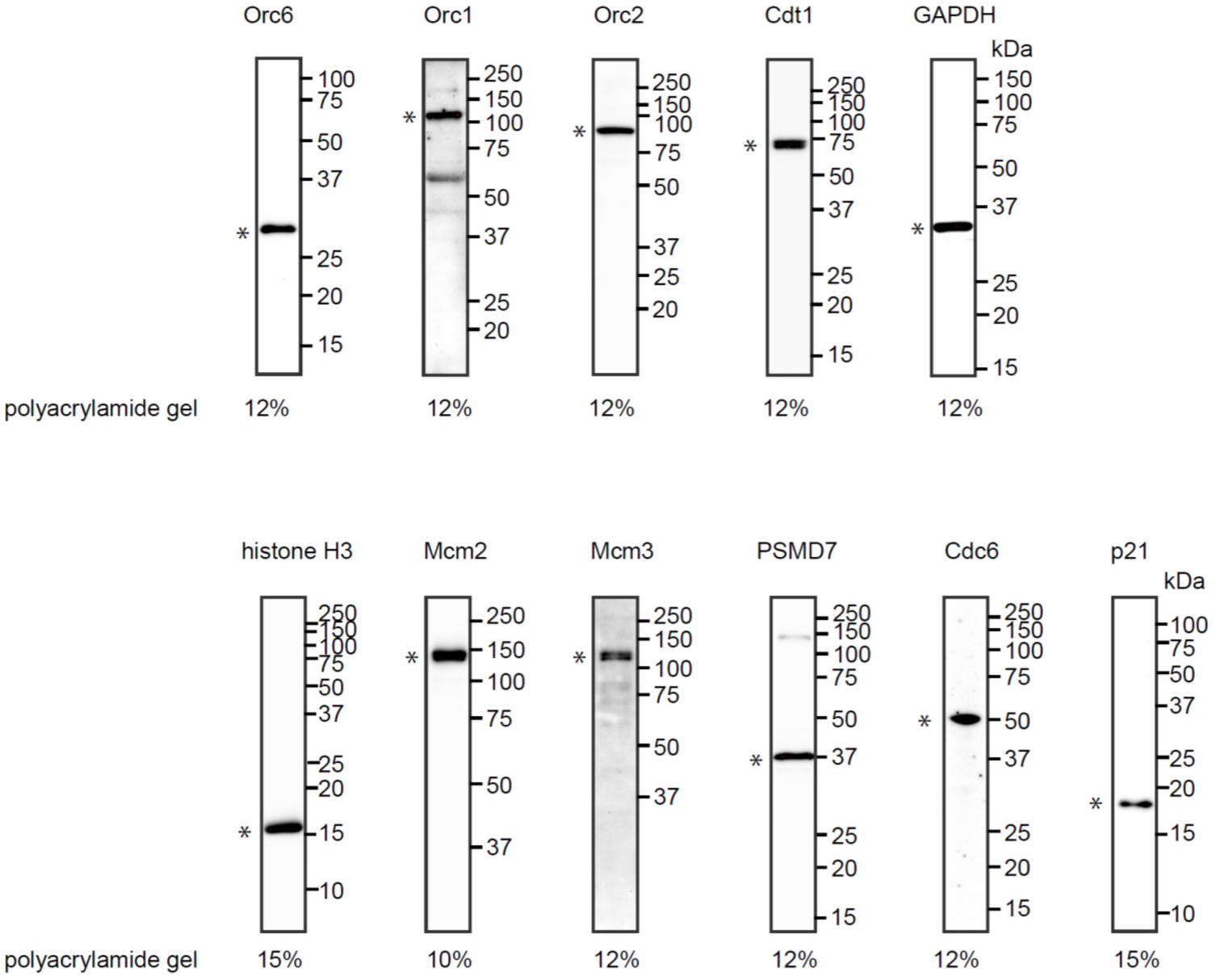
Validation of antibodies used in this study. Western blot analysis of whole-cell lysates from hTERT-RPE1 cells using antibodies against Orc6, Orc1, Orc2, Cdt1, GAPDH, histone H3, Mcm2, Mcm3, PSMD7, Cdc6, and p21. Molecular weight markers (kDa) are indicated on the right of each blot. The percentage of polyacrylamide gel used is shown below each blot. Asterisks (*) indicate the proteins of interest detected at their expected molecular weights.

**Figure S2.**
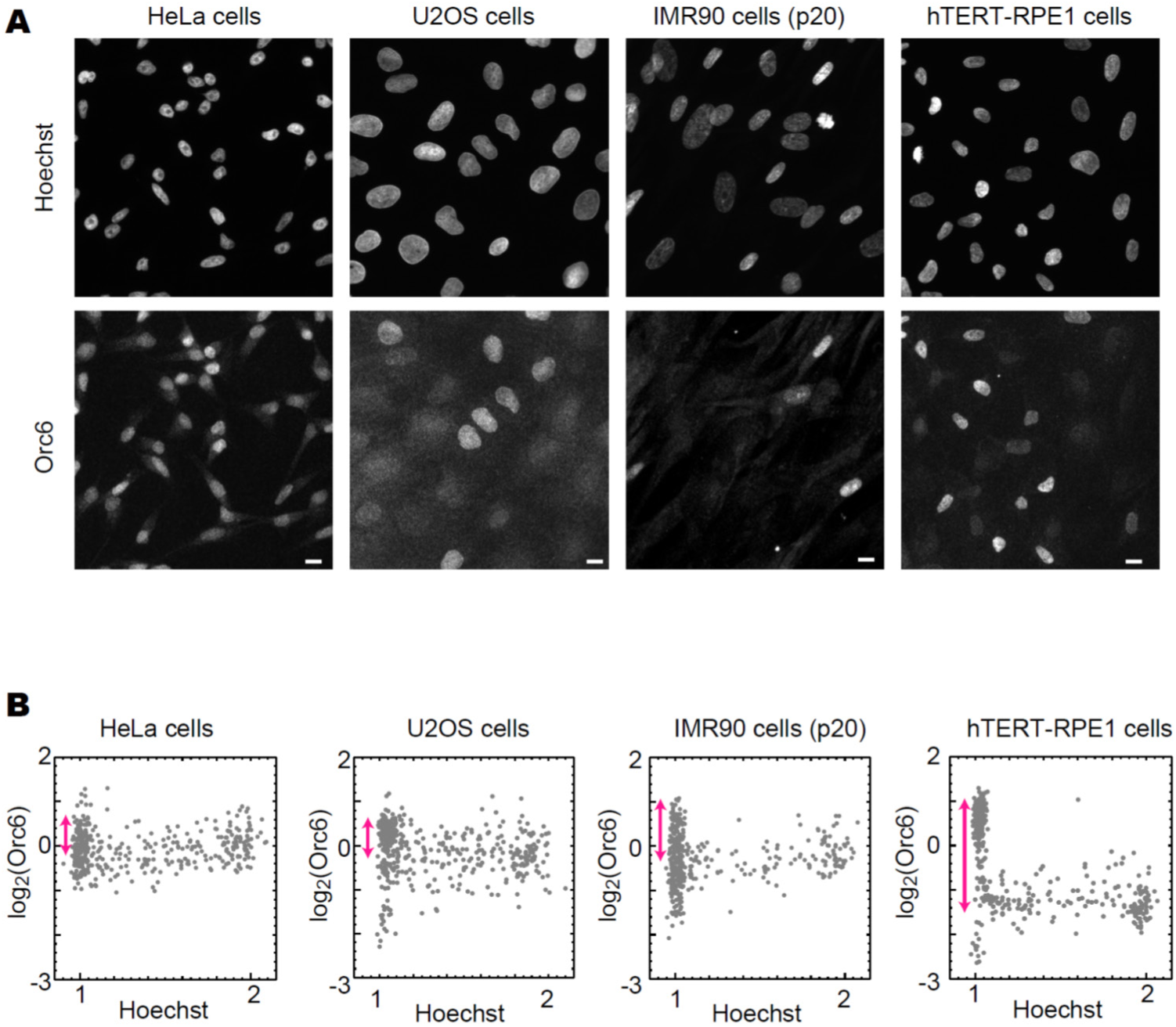
Behaviors of chromatin-bound Orc6 during the cell cycle in HeLa, U2OS, IMR90, and hTERT-RPE1 cells. (**A**) Representative fluorescence images of Hoechst and Orc6 staining in HeLa (left), U2OS (second left), IMR90 (second right), and hTERT-RPE1 (right) cells. Cells were fixed using the pre-extraction method and stained with Hoechst and anti-Orc6 antibody. The method of pre-extraction reproducibly makes the nuclei of HeLa cells smaller, but not in U2OS cells, for unknown reasons. Scale bars, 10 µm. (**B**) Single-cell plot analysis based on the images in (**A**). Each dot represents the intensities of Orc6 in an individual cell plotted against the Hoechst intensities. Magenta arrows show the difference in Orc6 levels from the 10^th^ highest intensity in G1 (Hoechst < 1.1) to the Orc6 average intensity in early S-phase (1.15 < Hoechst < 1.4). n = 450 (number of cells examined in each panel).

**Figure S3.**
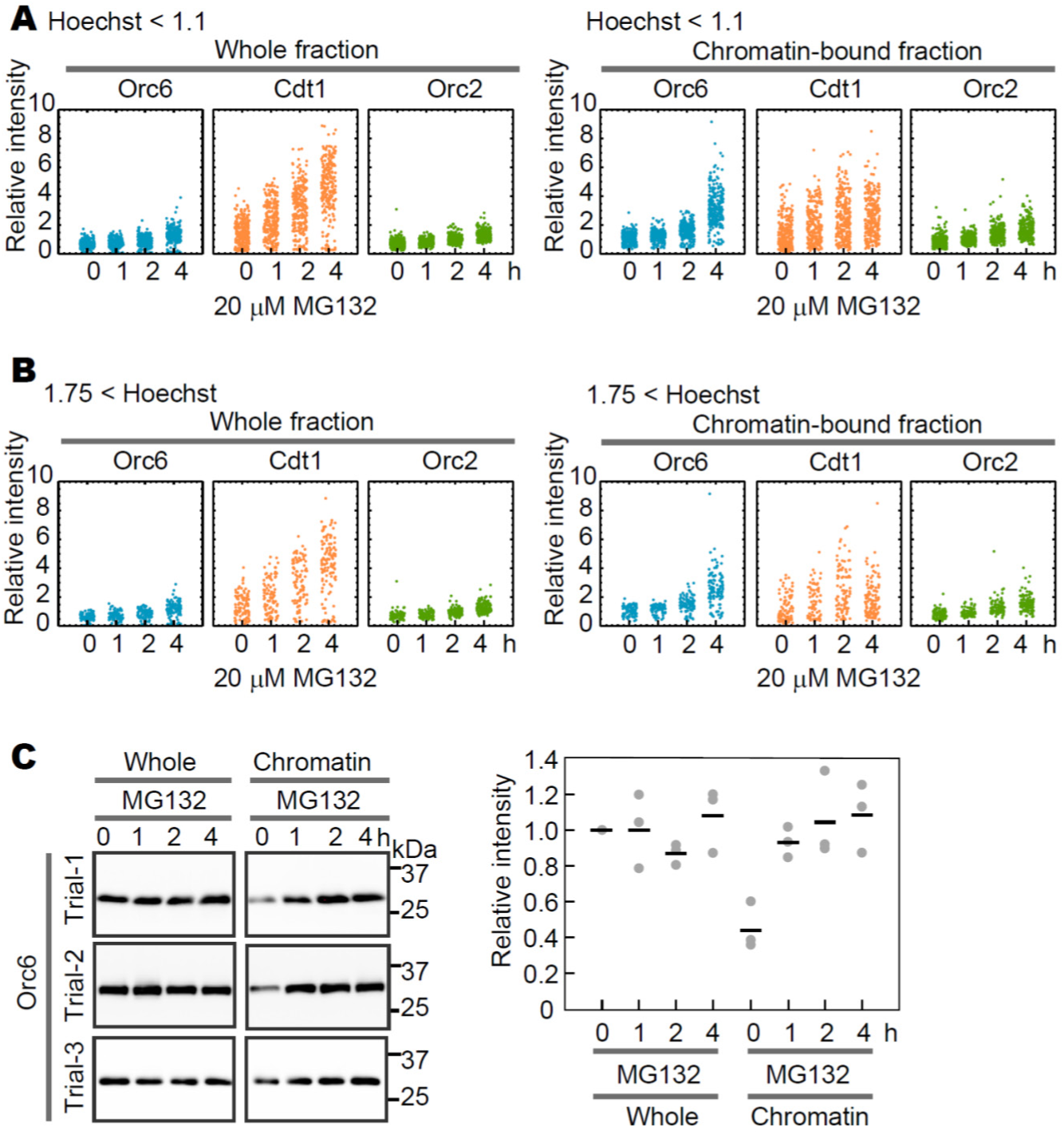
Chromatin binding of Orc6 is independent of the presence of Orc1 and Orc2. (**A, B**) Orc6, Cdt1, and Orc2 levels in G1-**(A)** and G2-phases cells **(B)**. Cells corresponding to G1-phase (Hoechst < 1.1) or G2-phase (1.75 < Hoechst) are selected from (Fig. 2A), and their fluorescence intensities were plotted. The average intensity in MG132-untreated cells (0 h) was defined as 1. **(C)** Western blot analysis using antibodies against Orc6. hTERT-RPE1 cells were treated with 20 μM MG132 for the indicated periods (h), as shown in Fig. 2D. MG132-treatment was carried out in three independent experiments, and analyzed by Western blot analysis for each experiment (left). The top panel is the same data shown in Fig. 2D. Band intensities were quantified by Image Lab software (BioRad) in each experiment, and normalized by the band intensity of the untreated whole-cell fraction (0 h, leftmost) set to 1. The normalized intensity of each band was plotted (right). The mean values from the three experiments are indicated by horizontal bars.

**Figure S4.**
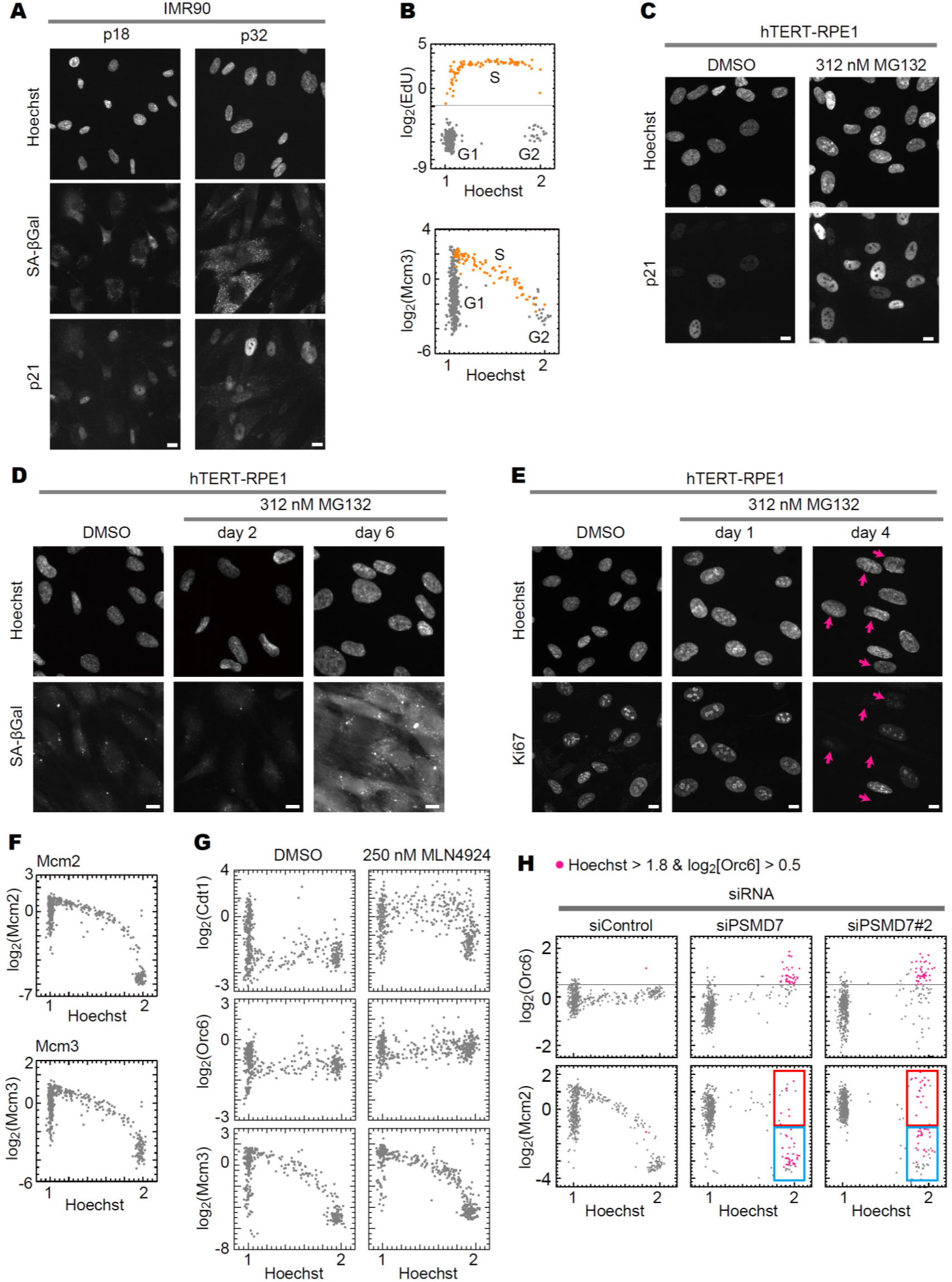
Effect of MG132 treatment on hTERT-RPE1 cells. (**A**) Representative fluorescence images of Hoechst, SA-βGal and p21 staining in IMR90 cells (passage numbers 18 and 32). Cells were fixed using the direct fixation method. **(B)** Single-cell plot analysis of IMR90 cells (passage 18) co-immunostained with EdU and Mcm3. Cells were treated with EdU for 30 min prior to collection and then fixed using the pre-extraction method. Each dot represents the intensities of EdU and Mcm3 in an individual cell, plotted against Hoechst 33342 intensities. Orange dots represent S-phase cells based on EdU intensities (log_2_(EdU) > ‒2). The number of cells examined in each panel is 450. (**C**) Representative fluorescence images of Hoechst and p21 staining in hTERT-RPE1 cells treated with DMSO or 312 nM MG132 on day 1. Cells were fixed using the direct fixation method. **(D)** Representative images of Hoechst and SA-βGal staining in cells treated with DMSO or 312 MG132 on day 2 and on day 6. Before the direct fixation method for Hoechst staining, cells were incubated with the reagent according to the manual. **(E)** Representative images of Hoechst and Ki67 staining in cells treated with DMSO or 312 nM MG132 on day 1 and on day 4. Cells were fixed using the direct fixation method. Red arrows indicate Ki67 negative cells. **(F)** Single-cell plot analysis of hTERT-RPE1 cells co-immunostained with Mcm2 and Mcm3. The number of cells examined in each panel is 450. (**G**) Single-cell plot analysis of hTERT-RPE1 cells treated with DMSO or 250 nM MLN4924 for 24 h. The cells were fixed using the pre-extraction method, and stained with Hoechst, anti-Orc6, anti-Mcm3, and anti-Cdt1 antibodies. n = 450 (number of cells examined in each panel). **(H)** Single-cell plot analysis of Orc6 and Mcm2 in hTERT-RPE1 cells treated with negative control-siRNA (siControl) or two types of PSMD7-siRNA (siPSMD7). Cell conditions and immunostaining methods were the same as in Fig.4E. n = 450 (number of cells examined in each panel). Scale bars, 10 µm.

**Figure S5.**
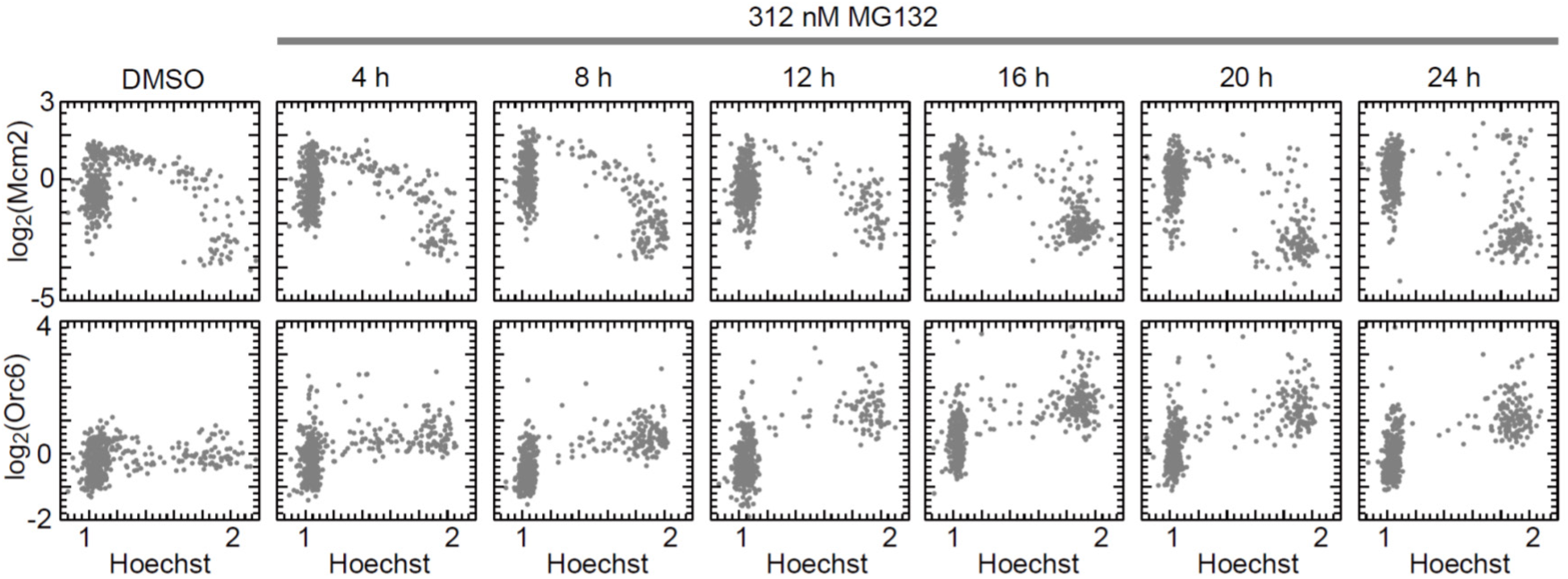
Time course from MG132 treatment to the appearance of cells with high chromatin-bound MCM and Orc6. Single-cell plot analysis of Mcm2 (top) and Orc6 (bottom) in cells treated with DMSO or 312 nM MG132 for the indicated durations. The same conditions but different combinations are shown in Fig. 5B, 5C. n = 450 (number of cells examined in each panel).

**Figure S6.**
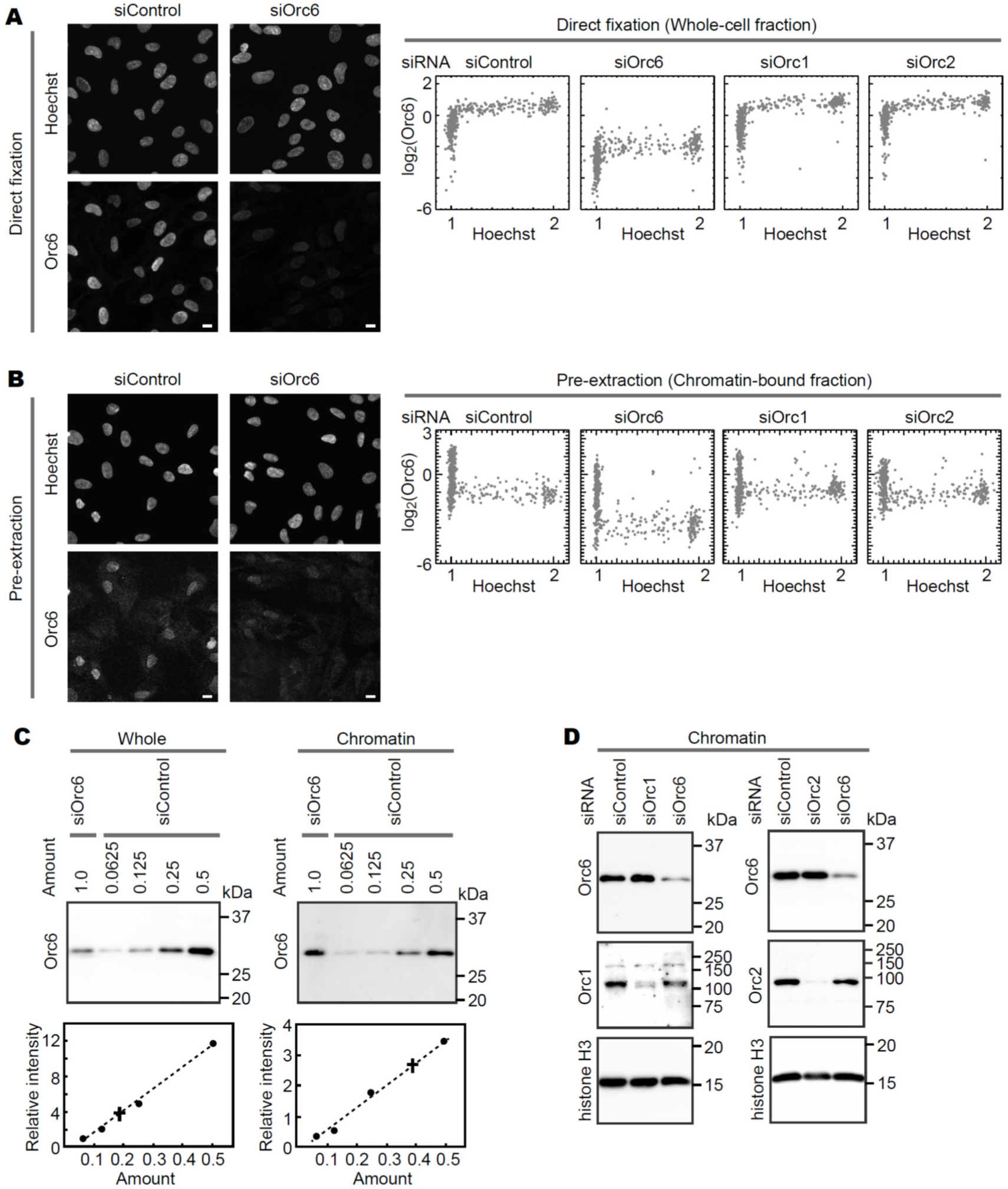
Effects of siRNA on Orc6 protein expression levels (A, **B)** Single-cell plot analysis of the Orc6 protein levels. The cells were treated with siRNA against a negative control (siControl), Orc6 (siOrc6), Orc1 (siOrc1), and Orc2 (siOrc2), and fixed using the direct fixation method to obtain the whole-cell fraction **(A)** or the pre-extraction method to obtaind the chromatin-bound fraction **(B)**. Left: Representative fluorescence images of Hoechst and Orc6 staining in hTERT-RPE1 cells for 24 h with siRNA against Orc6 (siOrc6) and a negative control (siControl). The images show that Orc6 levels are reduced by siOrc6 compared with siControl both in whole-cell fraction (direct fixation, A) and chromatin-bound fraction (pre-extraction, B). Scale bars, 10 µm. Right: Single-cell plot analysis based on these images for the cells treated with siRNA against a negative control (siControl), Orc6 (siOrc6), Orc1 (siOrc1), and Orc2 (siOrc2), and fixed using the direct fixation method **(A)** or the pre-extraction method **(B)**. n = 450 (number of cells examined in each panel). **(C)** Quantitative Western blot analysis of Orc6 siRNA. Cells were treated with siOrc6 or siControl. Chromatin-bound fractions were prepared from these cells using the sucrose cushion fractionation as described in Methods shown in Fig. 2C. Whole-cells (left) or chromatin-bound fractions (right) were electrophoresed, and analyzed by Western blot using Orc6 antibody (top). Amounts of siControl samples applied to each lane corresponded to 0.5, 0.25, 0.125, and 0.0625 relative to the amount of siOrc6 sample. Band intensities were quantified by Image Lab software (BioRad) in each experiment, and plotted to confirm linearity (bottom). The amounts of Orc6 in siOrc6-treated cells were estimated using a linear approximation curve (dash line) derived from siControl (solid circles); the plus symbol represents the level of siOrc6 equivalent to the linear regression of siControl (dash line). In cells treated with siORC6, the Orc6 levels in the fraction corresponding to 4 × 10^4^ cells (set as 1) were approximately 0.2 in the whole-cell fraction and 0.4 in the chromatin-bound fraction content compared to the control. (**D**) Western blot analysis using antibodies against Orc6, Orc1, Orc2, and histone H3. hTERT-RPE1 cells were treated with siRNA (siControl, siOrc1, siOrc2, or siOrc6) for 24 h. Chromatin-bound fractions were subjected to western blot analyses. For the Orc6 antibody staining, 2 × 10^4^ cells (equivalent to 0.5 in **C**) were applied to each lane. Histone H3 is a loading control. Molecular weight markers (kDa) are indicated on the right.

**Figure S7.**
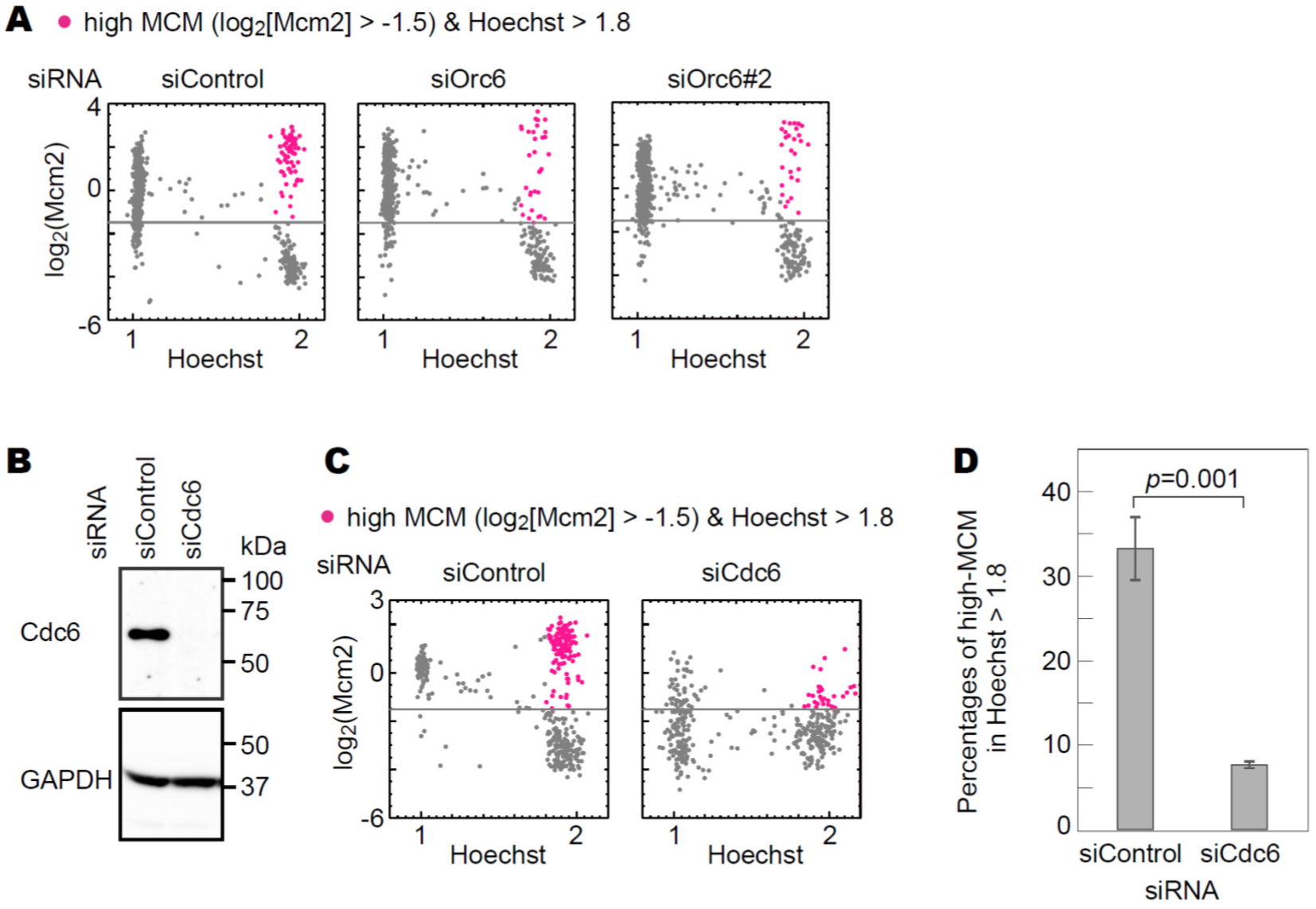
Single-cell plot analysis in siRNA-treated cells. **(A)** Single-cell plot analysis of Mcm2 in cells treated with siOrc6 and siOrc6#2. **(B)** Western blot analysis of whole-cell lysates from hTERT-RPE1 cells treated with siControl or siCdc6), using antibodies against Cdc6 and GAPDH. Molecular weight markers (kDa) are indicated on the right. (**C)** Single-cell plot analysis of chromatin-bound Mcm2 in hTERT-RPE1 cells treated with siControl or siCdc6. After MG132 treatment, cells were fixed using the pre-extraction method and stained with Hoechst and anti-Mcm2 antibody. Cells with high MCM levels (log_2_[Mcm2] > −1.5) and high DNA content (Hoechst > 1.8) are marked magenta. n = 450 (number of cells examined in each panel). (**D**) Percentage of magenta-marked cells (log_2_[Mcm2] > −1.5 and Hoechst > 1.8) among G2-phase cells (Hoechst > 1.8) based on (**C)**. Columns and error bars represent the mean and standard error of the mean of three independent experiments. Statistical difference (*p* = 0.001) was determined using a one-sided *t-*test.

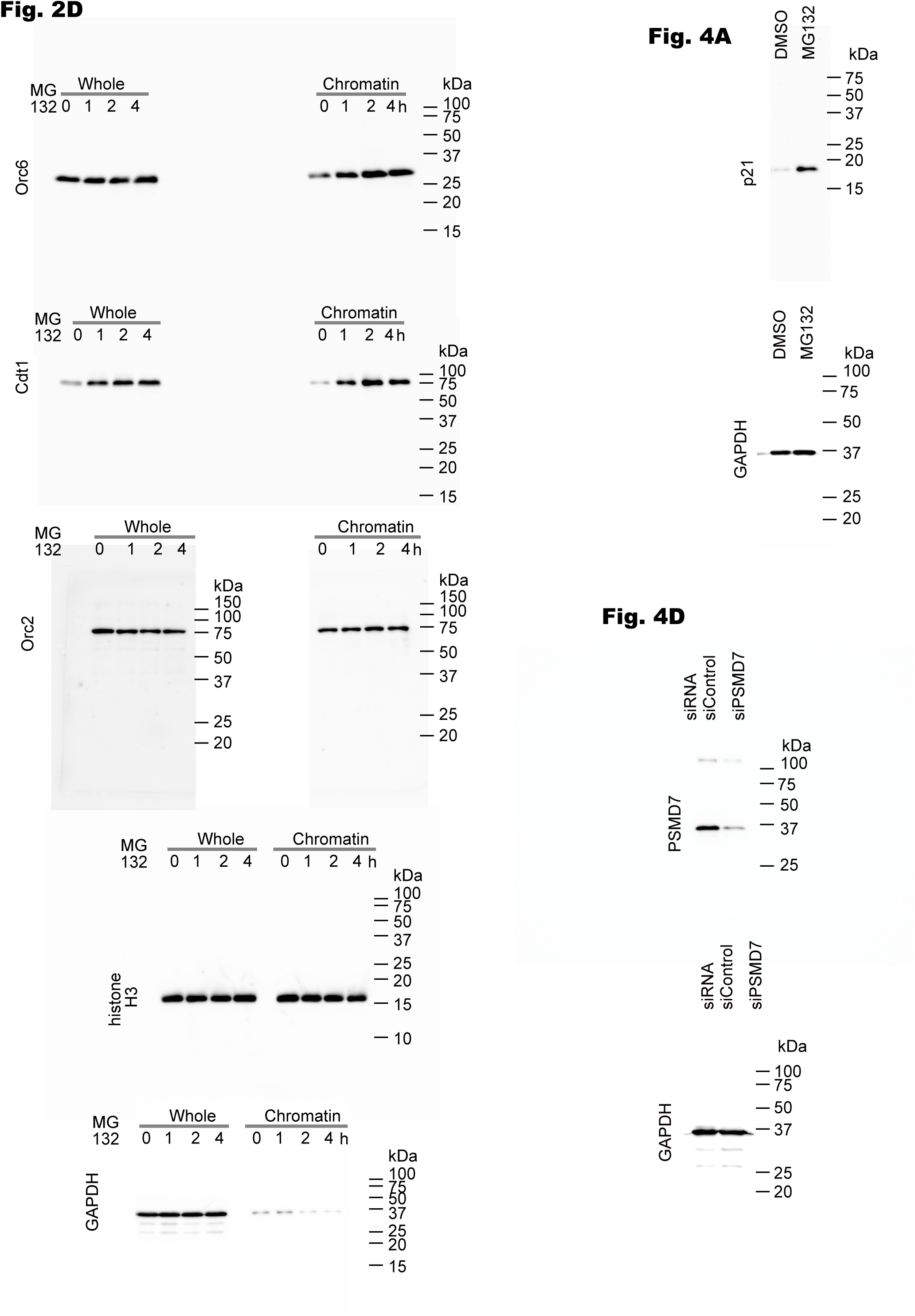

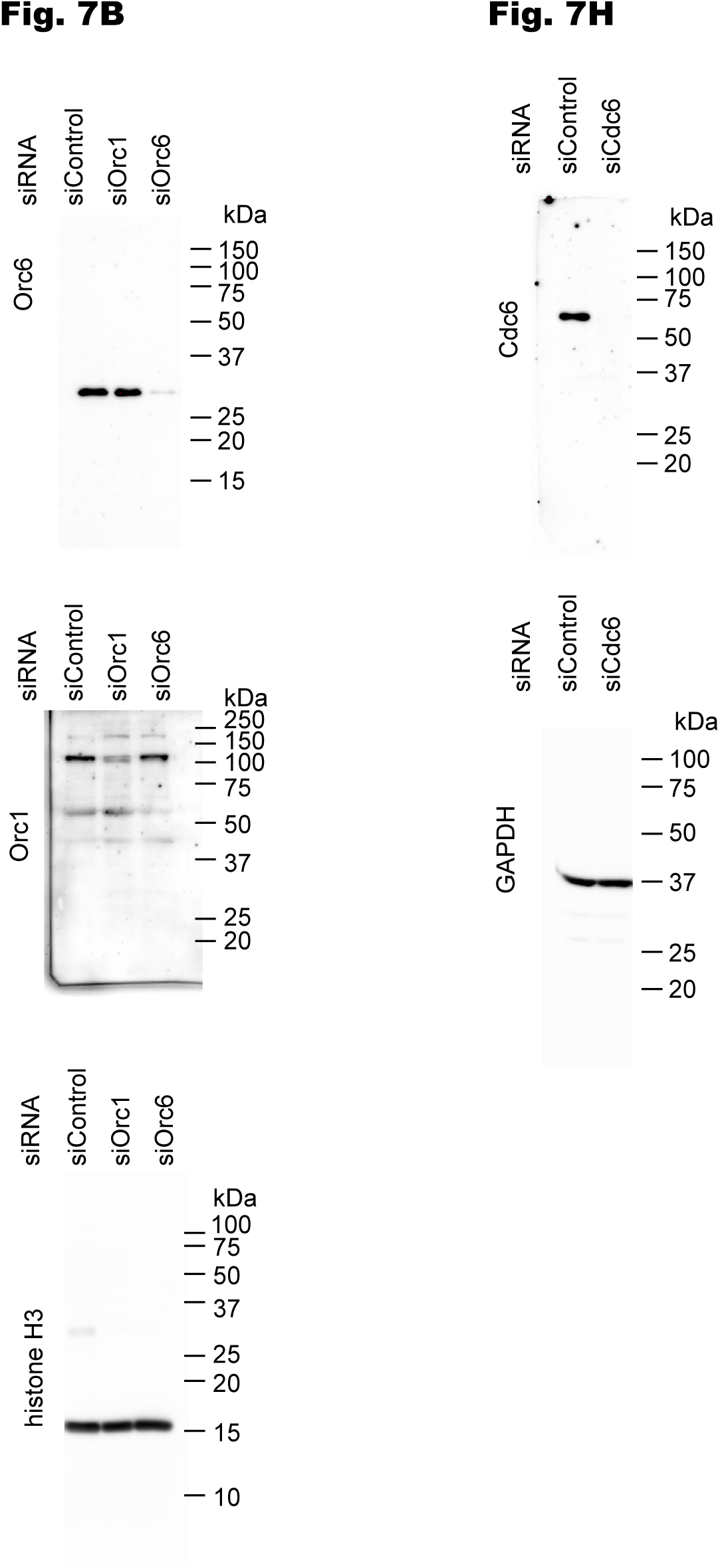

## References

Balasov, M., Huijbregts, R. P. H. and Chesnokov, I. (2009). Functional analysis of an Orc6 mutant in Drosophila. Proc Natl Acad Sci U S A 106, 10672–10677.

Bell, S. D. and Botchan, M. R. (2013). The minichromosome maintenance replicative helicase. Cold Spring Harb Perspect Biol 5, a012807.

Bell, S. P. and Labib, K. (2016). Chromosome Duplication in Saccharomyces cerevisiae. Genetics 203, 1027–1067.

Bernal, J. A. and Venkitaraman, A. R. (2011). A vertebrate N-end rule degron reveals that Orc6 is required in mitosis for daughter cell abscission. J Cell Biol 192, 969– 978.

Bielski, C. M., Zehir, A., Penson, A. V., Donoghue, M. T. A., Chatila, W., Armenia, J., Chang, M. T., Schram, A. M., Jonsson, P., Bandlamudi, C., et al. (2018). Genome doubling shapes the evolution and prognosis of advanced cancers. Nat Genet 50, 1189–1195.

Chen, S. and Bell, S. P. (2011). CDK prevents Mcm2–7 helicase loading by inhibiting Cdt1 interaction with Orc6. Genes Dev. 25, 363–372.

Chen, S., De Vries, M. A. and Bell, S. P. (2007). Orc6 is required for dynamic recruitment of Cdt1 during repeated Mcm2–7 loading. Genes Dev. 21, 2897–2907.

Chesnokov, I. N., Chesnokova, O. N. and Botchan, M. (2003). A cytokinetic function of *Drosophila* ORC6 protein resides in a domain distinct from its replication activity. Proc. Natl. Acad. Sci. U.S.A. 100, 9150–9155.

Chondrogianni, N. and Gonos, E. S. (2004). Proteasome inhibition induces a senescence-like phenotype in primary human fibroblasts cultures. Biogerontology 5, 55–61.

Chondrogianni, N., Stratford, F. L. L., Trougakos, I. P., Friguet, B., Rivett, A. J. and Gonos, E. S. (2003). Central role of the proteasome in senescence and survival of human fibroblasts: induction of a senescence-like phenotype upon its inhibition and resistance to stress upon its activation. J Biol Chem 278, 28026–28037.

Coulombe, P., Nassar, J., Peiffer, I., Stanojcic, S., Sterkers, Y., Delamarre, A., Bocquet, S. and Méchali, M. (2019). The ORC ubiquitin ligase OBI1 promotes DNA replication origin firing. Nat Commun 10, 2426.

Deegan, T. D. and Diffley, J. F. (2016). MCM: one ring to rule them all. Current Opinion in Structural Biology 37, 145–151.

Dhar, S. K. and Dutta, A. (2000). Identification and Characterization of the Human ORC6 Homolog. Journal of Biological Chemistry 275, 34983–34988.

Duncker, B. P., Chesnokov, I. N. and McConkey, B. J. (2009). The origin recognition complex protein family. Genome Biol 10, 214.

Feng, X., Noguchi, Y., Barbon, M., Stillman, B., Speck, C. and Li, H. (2021). The structure of ORC-Cdc6 on an origin DNA reveals the mechanism of ORC activation by the replication initiator Cdc6. Nat Commun 12, 3883.

Ganier, O., Prorok, P., Akerman, I. and Méchali, M. (2019). Metazoan DNA replication origins. Curr Opin Cell Biol 58, 134–141.

Gemble, S., Wardenaar, R., Keuper, K., Srivastava, N., Nano, M., Macé, A.-S., Tijhuis, A. E., Bernhard, S. V., Spierings, D. C. J., Simon, A., et al. (2022). Genetic instability from a single S phase after whole-genome duplication. Nature 604, 146–151.

Håland, T. W., Boye, E., Stokke, T., Grallert, B. and Syljuåsen, R. G. (2015). Simultaneous measurement of passage through the restriction point and MCM loading in single cells. Nucleic Acids Res 43, e150–e150.

Hayashi-Takanaka, Y., Yamagata, K., Nozaki, N. and Kimura, H. (2009). Visualizing histone modifications in living cells: spatiotemporal dynamics of H3 phosphorylation during interphase. J Cell Biol 187, 781–790.

Hayashi-Takanaka, Y., Yamagata, K., Wakayama, T., Stasevich, T. J., Kainuma, T., Tsurimoto, T., Tachibana, M., Shinkai, Y., Kurumizaka, H., Nozaki, N., et al. (2011). Tracking epigenetic histone modifications in single cells using Fab-based live endogenous modification labeling. Nucleic Acids Research 39, 6475–6488.

Hayashi-Takanaka, Y., Maehara, K., Harada, A., Umehara, T., Yokoyama, S., Obuse, C., Ohkawa, Y., Nozaki, N. and Kimura, H. (2015). Distribution of histone H4 modifications as revealed by a panel of specific monoclonal antibodies. Chromosome Res 23, 753–766.

Hayashi-Takanaka, Y., Kina, Y., Nakamura, F., Becking, L. E., Nakao, Y., Nagase, T., Nozaki, N. and Kimura, H. (2020). Histone modification dynamics as revealed by multicolor immunofluorescence-based single-cell analysis. J Cell Sci 133, jcs243444.

Hayashi-Takanaka, Y., Hayashi, Y., Hirano, Y., Miyawaki-Kuwakado, A., Ohkawa, Y., Obuse, C., Kimura, H., Haraguchi, T. and Hiraoka, Y. (2021). Chromatin loading of MCM hexamers is associated with di-/tri-methylation of histone H4K20 toward S phase entry. Nucleic Acids Res 49, 12152–12166.

Hu, Y. and Stillman, B. (2023). Origins of DNA replication in eukaryotes. Mol Cell 83, 352–372.

Huang, W., Hickson, L. J., Eirin, A., Kirkland, J. L. and Lerman, L. O. (2022). Cellular senescence: the good, the bad and the unknown. Nat Rev Nephrol 18, 611–627.

Ishimi, Y. (2018). Regulation of MCM2-7 function. Genes Genet Syst 93, 125–133.

Johmura, Y., Shimada, M., Misaki, T., Naiki-Ito, A., Miyoshi, H., Motoyama, N., Ohtani, N., Hara, E., Nakamura, M., Morita, A., et al. (2014). Necessary and Sufficient Role for a Mitosis Skip in Senescence Induction. Molecular Cell 55, 73– 84.

Kinyamu, H. K., Bennett, B. D., Bushel, P. R. and Archer, T. K. (2020). Proteasome inhibition creates a chromatin landscape favorable to RNA Pol II processivity. J Biol Chem 295, 1271–1287.

Kwon, Y. H., Jovanovic, A., Serfas, M. S., Kiyokawa, H. and Tyner, A. L. (2002). P21 functions to maintain quiescence of p27-deficient hepatocytes. J Biol Chem 277, 41417–41422.

Li, C.-J. and DePamphilis, M. L. (2002). Mammalian Orc1 Protein Is Selectively Released from Chromatin and Ubiquitinated during the S-to-M Transition in the Cell Division Cycle. Molecular and Cellular Biology 22, 105–116.

Limas, J. C. and Cook, J. G. (2019). Preparation for DNA replication: the key to a successful S phase. FEBS Lett 593, 2853–2867.

Lin, J. J., Milhollen, M. A., Smith, P. G., Narayanan, U. and Dutta, A. (2010). NEDD8-targeting drug MLN4924 elicits DNA rereplication by stabilizing Cdt1 in S phase, triggering checkpoint activation, apoptosis, and senescence in cancer cells. Cancer Res 70, 10310–10320.

Lin, Y.-C., Liu, D., Chakraborty, A., Kadyrova, L. Y., Song, Y. J., Hao, Q., Mitra, J., Hsu, R. Y. C., Arif, M. K., Adusumilli, S., et al. (2022). Orc6 is a component of the replication fork and enables efficient mismatch repair. Proc Natl Acad Sci U S A 119, e2121406119.

Lin, Y.-C., Liu, D., Chakraborty, A., Macias, V., Brister, E., Sonalkar, J., Shen, L., Mitra, J., Ha, T., Kajdacsy-Balla, A., et al. (2023). DNA Damage-Induced, S-Phase Specific Phosphorylation of Orc6 is Critical for the Maintenance of Genome Stability. Mol Cell Biol 43, 143–156.

Liu, S., Balasov, M., Wang, H., Wu, L., Chesnokov, I. N. and Liu, Y. (2011). Structural analysis of human Orc6 protein reveals a homology with transcription factor TFIIB. Proc. Natl. Acad. Sci. U.S.A. 108, 7373–7378.

López, S., Lim, E. L., Horswell, S., Haase, K., Huebner, A., Dietzen, M., Mourikis, T. P., Watkins, T. B. K., Rowan, A., Dewhurst, S. M., et al. (2020). Interplay between whole-genome doubling and the accumulation of deleterious alterations in cancer evolution. Nat Genet 52, 283–293.

Makino, Y., Yogosawa, S., Kanemaki, M., Yoshida, T., Yamano, K., Kishimoto, T., Moncollin, V., Egly, J. M., Muramatsu, M. and Tamura, T. (1996). Structures of the rat proteasomal ATPases: determination of highly conserved structural motifs and rules for their spacing. Biochem Biophys Res Commun 220, 1049– 1054.

Makino, Y., Yoshida, T., Yogosawa, S., Tanaka, K., Muramatsu, M. and Tamura, T. A. (1999). Multiple mammalian proteasomal ATPases, but not proteasome itself, are associated with TATA-binding protein and a novel transcriptional activator, TIP120. Genes Cells 4, 529–539.

Méndez, J., Zou-Yang, X. H., Kim, S.-Y., Hidaka, M., Tansey, W. P. and Stillman, B. (2002). Human Origin Recognition Complex Large Subunit Is Degraded by Ubiquitin-Mediated Proteolysis after Initiation of DNA Replication. Molecular Cell 9, 481–491.

Nishitani, H., Sugimoto, N., Roukos, V., Nakanishi, Y., Saijo, M., Obuse, C., Tsurimoto, T., Nakayama, K. I., Nakayama, K., Fujita, M., et al. (2006). Two E3 ubiquitin ligases, SCF-Skp2 and DDB1-Cul4, target human Cdt1 for proteolysis. EMBO J 25, 1126–1136.

Nozawa, R.-S., Nagao, K., Masuda, H.-T., Iwasaki, O., Hirota, T., Nozaki, N., Kimura, H. and Obuse, C. (2010). Human POGZ modulates dissociation of HP1alpha from mitotic chromosome arms through Aurora B activation. Nat Cell Biol 12, 719– 727.

Ogawa, Y., Takahashi, T. and Masukata, H. (1999). Association of fission yeast Orp1 and Mcm6 proteins with chromosomal replication origins. Mol Cell Biol 19, 7228– 7236.

Pan, Q., Li, F., Ding, Y., Huang, H. and Guo, J. (2022). ORC6 acts as a biomarker and reflects poor outcome in clear cell renal cell carcinoma. J Cancer 13, 2504–2514.

Panopoulos, A., Pacios-Bras, C., Choi, J., Yenjerla, M., Sussman, M. A., Fotedar, R. and Margolis, R. L. (2014). Failure of cell cleavage induces senescence in tetraploid primary cells. Mol Biol Cell 25, 3105–3118.

Prasanth, S. G., Prasanth, K. V. and Stillman, B. (2002). Orc6 Involved in DNA Replication, Chromosome Segregation, and Cytokinesis. Science 297, 1026–1031.

Quan, Y., Xia, Y., Liu, L., Cui, J., Li, Z., Cao, Q., Chen, X. S., Campbell, J. L. and Lou, H. (2015). Cell-Cycle-Regulated Interaction between Mcm10 and Double Hexameric Mcm2-7 Is Required for Helicase Splitting and Activation during S Phase. Cell Reports 13, 2576–2586.

Sabath, N., Levy-Adam, F., Younis, A., Rozales, K., Meller, A., Hadar, S., Soueid-Baumgarten, S. and Shalgi, R. (2020). Cellular proteostasis decline in human senescence. Proc Natl Acad Sci U S A 117, 31902–31913.

Samson, R. Y., Abeyrathne, P. D. and Bell, S. D. (2016). Mechanism of Archaeal MCM Helicase Recruitment to DNA Replication Origins. Mol Cell 61, 287–296.

Scholzen, T. and Gerdes, J. (2000). The Ki-67 protein: from the known and the unknown. J Cell Physiol 182, 311–322.

Semple, J. W., Da-Silva, L. F., Jervis, E. J., Ah-Kee, J., Al-Attar, H., Kummer, L., Heikkila, J. J., Pasero, P. and Duncker, B. P. (2006). An essential role for Orc6 in DNA replication through maintenance of pre-replicative complexes. EMBO J 25, 5150–5158.

Sheu, Y.-J. and Stillman, B. (2006). Cdc7-Dbf4 Phosphorylates MCM Proteins via a Docking Site-Mediated Mechanism to Promote S Phase Progression. Molecular Cell 24, 101–113.

Shi, K., Zhang, J.-Z., Zhao, R.-L., Yang, L. and Guo, D. (2018). PSMD7 downregulation induces apoptosis and suppresses tumorigenesis of esophageal squamous cell carcinoma via the mTOR/p70S6K pathway. FEBS Open Bio 8, 533–543.

Soucy, T. A., Smith, P. G. and Rolfe, M. (2009). Targeting NEDD8-activated cullin-RING ligases for the treatment of cancer. Clin Cancer Res 15, 3912–3916.

Sugimoto, N., Maehara, K., Yoshida, K., Ohkawa, Y. and Fujita, M. (2018). Genome-wide analysis of the spatiotemporal regulation of firing and dormant replication origins in human cells. Nucleic Acids Res 46, 6683–6696.

Tang, M., Chen, J., Zeng, T., Ye, D.-M., Li, Y.-K., Zou, J. and Zhang, Y.-P. (2023). Systemic analysis of the DNA replication regulator origin recognition complex in lung adenocarcinomas identifies prognostic and expression significance. Cancer Med 12, 5035–5054.

Tatsumi, Y., Ohta, S., Kimura, H., Tsurimoto, T. and Obuse, C. (2003). The ORC1 Cycle in Human Cells. Journal of Biological Chemistry 278, 41528–41534.

Thakur, B. L., Ray, A., Redon, C. E. and Aladjem, M. I. (2022). Preventing excess replication origin activation to ensure genome stability. Trends Genet 38, 169– 181.

Torres, C., Lewis, L. and Cristofalo, V. J. (2006). Proteasome inhibitors shorten replicative life span and induce a senescent-like phenotype of human fibroblasts. J Cell Physiol 207, 845–853.

Ukekawa, R., Maegawa, N., Mizutani, E., Fujii, M. and Ayusawa, D. (2004). Proteasome inhibitors induce changes in chromatin structure characteristic of senescent human fibroblasts. Biosci Biotechnol Biochem 68, 2395–2397.

Vassilev, A. and DePamphilis, M. (2017). Links between DNA Replication, Stem Cells and Cancer. Genes 8, 45.

Vilchez, D., Saez, I. and Dillin, A. (2014). The role of protein clearance mechanisms in organismal ageing and age-related diseases. Nat Commun 5, 5659.

Walter, D., Hoffmann, S., Komseli, E.-S., Rappsilber, J., Gorgoulis, V. and Sørensen, C. S. (2016). SCFCyclin F-dependent degradation of CDC6 suppresses DNA re-replication. Nat Commun 7, 10530.

Xu, N., You, Y., Liu, C., Balasov, M., Lun, L. T., Geng, Y., Fung, C. P., Miao, H., Tian, H., Choy, T. T., et al. (2020). Structural basis of DNA replication origin recognition by human Orc6 protein binding with DNA. Nucleic Acids Research 48, 11146–11161.

Zeng, J., Hills, S. A., Ozono, E. and Diffley, J. F. X. (2023). Cyclin E-induced replicative stress drives p53-dependent whole-genome duplication. Cell 186, 528–542.e14.

Zhou, Y., Pozo, P. N., Oh, S., Stone, H. M. and Cook, J. G. (2020). Distinct and sequential re-replication barriers ensure precise genome duplication. PLoS Genet 16, e1008988.

Zhu, Q., Wani, G., Yao, J., Patnaik, S., Wang, Q.-E., El-Mahdy, M. A., Praetorius-Ibba, M. and Wani, A. A. (2007). The ubiquitin-proteasome system regulates p53-mediated transcription at p21waf1 promoter. Oncogene 26, 4199–4208.

